# Mass Spectrometry-Based Profiling of Personalized Immunopeptidomes in Thai Renal Cell Carcinoma

**DOI:** 10.64898/2026.01.03.696780

**Authors:** Poorichaya Somparn, Sira Sriswasdi, Piriya Wongkongkathep, Jirapat Techachakrit, Phijitra Muanwien, Saharat Nanthawong, Tatchapon Apinan, Julin Opanuraks, Shanop Shuangshoti, Nattiya Hirankarn, Sutpirat Moonmuang, Tanyaluck Kampoun, Parunya Chaiyawat, Trairak Pisitkun

**Author notes:** Corresponding authors: Correspondence to Trairak Pisitkun.

## Abstract

This study profiles the personalized immunopeptidomes of 13 Thai patients with renal cell carcinoma (RCC), addressing a critical knowledge gap in Southeast Asian populations characterized by distinct HLA allele distributions. We combined whole-exome sequencing (WES)–based personalized proteome construction with liquid chromatography–tandem mass spectrometry (LC–MS/MS), using both database-driven searches and *de novo* peptide sequencing. HLA typing identified seven alleles not previously represented in major immunopeptidome databases, with HLA-A*11:01 being the most frequent (69%). Database-based analysis identified a single tumor-specific neoantigen derived from a mutant JADE2 peptide in the patient with the highest tumor mutational burden, which was validated by a mutant-specific ELISPOT response. In contrast, *de novo* sequencing revealed numerous non-canonical peptides, a subset of which were supported by proteogenomic validation using PepQuery and detected exclusively in cancer proteomes but not in normal tissue datasets, indicating their potential as tumor-associated antigens. Together, these results establish an integrated and scalable framework for identifying HLA-presented tumor-derived peptides and provide a foundational immunopeptidome resource to support personalized cancer immunotherapy development in Southeast Asia.

## INTRODUCTION

Renal cell carcinoma (RCC) represents a significant global health burden, constituting a major portion of kidney cancer diagnoses and contributing considerably to cancer-related mortality [1]. Despite the transformative impact of immune checkpoint blockade (ICB) therapies in RCC treatment, their clinical efficacy remains inconsistent [2], highlighting the urgent need for personalized therapeutic strategies tailored to the unique immunological context of each patient. Promising advances in personalized immunotherapy, such as neoantigen vaccines, have demonstrated encouraging results in various cancer types, including melanoma, glioblastoma, and lung cancer [3-5]. The immunopeptidome—the ensemble of peptides presented by human leukocyte antigen (HLA) molecules on the surface of tumor cells—is central to advancing such personalized approaches [6]. These peptides, which are often derived from tumor-associated or mutation-derived proteins, have the potential to be recognized by T cells and trigger a robust anti-tumor immune response.

Mass spectrometry-based immunopeptidomics has emerged as an essential technology for the high-throughput and sensitive identification of these potential therapeutic targets [7]. Detailed immunopeptidome characterization enables the identification of tumor-derived antigens, including neoantigens, which are typically unique to an individual’s tumor and can serve as prime targets for personalized immunotherapy. The ethnic diversity of HLA alleles presents profound clinical implications, as variations across populations can significantly influence the immunopeptidome, ultimately affecting tumor antigen presentation and the effectiveness of immunotherapies [8]. Nonetheless, most immunopeptidome studies have been predominantly focused on Western populations, resulting in a considerable knowledge gap regarding diverse immunopeptidome landscapes [9-13]. The HLA Ligand Atlas database reveals a predominance of publications from Western countries, with 46 studies originating from the USA, Switzerland, Germany, and England, while only 2 studies have been conducted in Japan, and none from Southeast Asian countries [9]. This gap underscores the necessity for targeted research on underrepresented populations, particularly those in Southeast Asia. A lack of insight into the distinct HLA allele distributions in non-Western populations hampers the optimization of immunotherapies, potentially limiting their success and contributing to disparities in healthcare outcomes for these groups. Southeast Asian populations, including the Thai population, exhibit unique HLA allele distributions that may significantly impact immunotherapy outcomes. Thailand’s distinct HLA allele patterns and inherent genetic diversity offer a valuable opportunity to explore and characterize the broader Southeast Asian immunopeptidome landscape [14].

Given the limited representation of Southeast Asian populations in immunopeptidome research, we employed state-of-the-art mass spectrometry techniques to comprehensively profile the immunopeptidomes of Thai RCC patients, aiming to develop tailored immunotherapy strategies for this region. Traditional immunopeptidome database search methods primarily rely on matching MS/MS spectra to known peptide sequences in reference databases [15]. However, such methods are limited in their ability to detect tumor antigens arising from unique genomic aberrations present in individual patient tumors. To address this limitation, we employed advanced computational methods, including personalized proteome database construction based on somatic mutations identified through whole-exome sequencing of each patient, coupled with *de novo* sequencing to determine peptide sequences directly from MS/MS spectra without relying on existing databases. Our analysis revealed numerous immunopeptidome sequences that align with known motifs of Southeast Asian HLA alleles. This research has significant implications not only for the Thai population but for Southeast Asia as a whole. By establishing a comprehensive immunopeptidome landscape for Thai RCC patients, this study contributes a reference immunopeptidome resource for the region, facilitating future research and supporting the development of personalized cancer immunotherapy.

## METHODS

### Patient selection

All patients diagnosed with RCC by clinical and radiological findings who underwent nephrectomy in the King Memorial Chulalongkorn Hospital (KCMH) between June 2020 to June 2021 were consecutively enrolled. According to the final pathological diagnosis, patients with histological subtypes other than RCC were excluded. Eventually, the absolute number of patients participating in this study was 13 cases. The Institutional Review Board of the Faculty of Medicine, Chulalongkorn University, Bangkok, Thailand has approved the present study which is to be carried out in compliance with the international guidelines for human research protection as Declaration of Helsinki, The Belmont Report, CIOMS Guideline and International Conference on Harmonization in Good Clinical Practice (ICH-GCP), IRB No. 411/63 and COA No. 752/2020. All written informed consents were acquired from all enrolled patients.

### Whole exome sequencing

Genomic DNA was extracted from snap-frozen tumor tissue and peripheral blood mononuclear cells (PBMCs) using the QIAGEN AllPrep kit. DNA libraries were prepared using the SureSelect Human All Exon V7 kit (Agilent USA) and sequenced on an MGI DNBSEQ-G400 platform (MGI Tech Co., Ltd, China) using a 2×150 bp paired-end configuration. The targeted average depth was at least 300x for tumor DNA and 50x for normal DNA. The quality of raw reads was assessed by using FastQC to analyze the GC content and quality scores [16]. After removing duplicates, the processed reads were aligned with reference genome version hg38 using Burrows-Wheeler Aligner (BWA version 0.7.15) [17] and further processed with the Genome Analysis Toolkit (GATK version 3.6) [17]. The aligned reads were then stored in BAM file (.bam). For the assembled genome data, the Picard tool was combined with Bamtools to filter out the mismatching and low-quality reads. The data distribution and reads coverage were then evaluated using the CalculateHsMetrics package.

### Variant identification and HLA typing

To identify somatic mutations, MuTect2 (version 4.1.4.0) [18], VarScan (version 2.4.2) [19], and Strelka (version 2.9.9) [20] were used. HLA allotypes were identified by Optitype [21] and HLA-LA [22]. In addition, TMB was determined as the number of all SNV mutations and indels per megabase of the genome examined [23].

### Pipeline for customized protein database

For each patient, the list of identified somatic and germline mutations in the coding sequences was used to create a personalized protein database for subsequent mass spectra search. Somatic mutations identified from the three tools were aggregated. Given the lack of haplotype information, peptides containing all possible combinations of somatic and germline mutations that occur within a window of 15 amino acid residues (45 nucleotide positions on the coding sequence) were generated and appended to the UniProt human reference proteome to derive the personalized protein database. The choice of 15 amino acid window was selected because it is the maximum length of HLA class I antigens assumed by downstream bioinformatics tools.

### In-house purification of antibody

Purified pan HLA-A, -B, -C was generated from W6/32 hybridoma cells (a gift from Noriko Steiner) cultured in DMEM media supplemented with 10% fetal bovine serum, L-glutamine, L glucose, and Sodium Pyruvate and 50 U/mL penicillin and expanded in roller bottles at 37 °C with 5% CO_2_. Secreted monoclonal antibodies were harvested from spent media and purified using Protein A resin with ÄKTA purification system (Cytiva, USA).

### The IP-beads preparation and MHC immunoprecipitation

Ten micrograms of W6/32 antibody was cross linked onto 1 ml of slurry of Protein G Sepharose (GE) by DMP (20 mM) in 0.2 M Triethanolamine pH 8.2 for 1 hour. After that washing with 10 ml of ice-cold 0.2 M Tris (pH 8), 10 ml 0.1 M Citrate buffer (pH3) and 10 ml of PBS, respectively, and stored at 4 °C until use. Tissue samples were added to 1× lysis buffer and pulverized using an MM400 Retsch Mixer Mill (Retsch, Germany), then lysed with 0.1% IGEPAL CA-630, 100 mM Tris, 300 mM NaCl, pH 8.0 Complete Protease Inhibitor Cocktail (Roche, Switzerland). The supernatant was passed through a Protein G resin pre-column (500 μL) to remove non-specific binding materials. HLA class I and II immunoaffinity purification was performed as previously described [24]. Briefly, the pre-cleared supernatant was incubated with 10 mg of pan HLA-A, -B, and -C antibodies coupled to Protein G resin with rotation overnight at 4 °C. After conjugation, the resins were washed with 10 ml of ice-cold wash buffer 1 (0.005% IGEPAL, 50 mM Tris, pH 8.0, 150 mM NaCl, 5 mM EDTA), 10 ml of ice-cold wash buffer 2 (50 mM Tris, pH 8.0, 150 mM NaCl), and 10 ml of ice-cold wash buffer 3 (50 mM Tris, pH 8.0, 450 mM NaCl). Bound complexes were eluted from the column using 5 column volumes of 10% acetic acid. Eluted peptides were fractionated by reverse-phase high-performance liquid chromatography (Shimadzu, Japan) on a 4.6 mm diameter Chromolith SpeedROD RP-18 (Merck, USA). The optimized conditions were as follows: mobile phase A (0.05% v/v TFA, 2.5% v/v ACN in water), mobile phase B (0.045% v/v TFA, 90% v/v ACN in water), flow rate of 1 mL/minute, temperature of 30 °C, and injection volume of 200 μL. The elution program was set as follows: 0-5% of mobile phase B over 1 minute, 5-15% of mobile phase B over 4 minutes, 15-45% of mobile phase B over 30 minutes, 45-100% of mobile phase B over 15 minutes, and 100% of mobile phase B over 4 minutes. Fractions were collected in 1 mL each. Consecutive fractions were pooled into 11 fractions. Pooled fractions were concentrated by vacuum centrifugation and reconstituted in 0.1% FA.

### LC-MS/MS Analysis

Pooled peptide fractions eluted from an HLA class I sample were analyzed on a Q Exactive mass spectrometer (Thermo Fisher Scientific, USA) coupled to an EASY-nLC 1000 (Thermo Fisher Scientific, USA). Peptide samples were separated at a flow rate of 300 nL /minute of buffer B (80% ACN, 0.1% FA). The gradient was set at 4-20% of buffer B over 30 minutes, 20-28% of buffer B over 40 minutes, 28-40% of buffer B over 5 minutes, 40-95% of buffer B over 3 minutes, washing with 95% of buffer B over 8 minutes, re-equilibration with buffer A (2% ACN/0.1% FA) over 5 minutes. Mass spectra resolutions were set at 70,000 for full MS scans and 17,500 for MS/MS scans. The normalized collision energy for HCD fragmentation was set at 30%. The m/z scan range was set at 350-1,400. Dynamic exclusion was set at 15 seconds. The maximum injection times were set at 120 ms for full MS scan and 120 ms for MS/MS scans. Each sample was analyzed twice on the mass spectrometer, once where precursor charge states of +2 or higher were accepted (raw file names beginning with 2zup) and another where all charge states were accepted (raw file names beginning with 1zup).

### Peptide Identification

The raw files were searched against the personalized protein databases constructed from the UniProt reference human proteome and the mutation profiles of individual patients using PEAKS Xpro and SMSnet *denovo* sequencing software to obtain peptide sequence information. For PEAKS Xpro, no enzyme was specified for enzymatic digestion. The precursor mass tolerance was set to 10 ppm and the fragment mass tolerance to 0.02 Da. The maximum number of missed cleavages was set to zero. Methionine oxidation and phosphorylation were included as dynamic modifications. A false discovery rate (FDR) was set at 0.05. Data have been deposited in the PRIDE Archive [25] at www.ebi.ac.uk/pride/archive (PXD072437). For SMSnet denovo sequencing, MS/MS Raw files were converted to Mascot Generic Format (MGF) using MSConvert from the ProteoWizard suite [26]. Vendor-specific peak picking algorithms were employed for both MS1 and MS2 spectra. Zero-intensity samples were removed from MS2 spectra, and only MS2 spectra were converted. The default title maker was utilized for file naming. Charge state deconvolution was not performed during the conversion process. The SMSNet-M model which treats carbamidomethylation of cysteine as fixed modification and oxidation of methionine as variable modification was used [27]. Target amino acid-level false discovery rate was set at 5%. Precursor mass tolerance of 30 ppm was applied to discard identified peptides with high mass deviations. Partially identified peptides were searched against a customized database combined reference human proteome form UniProt.

### Peptide clustering analysis and Peptide-HLA binding prediction by GibbCluster and NetMHCpan-4.1

We performed peptide clustering analysis and peptide-HLA binding prediction using GibbsCluster for peptide sequence alignment and clustering, providing a comprehensive and unbiased view of collective motifs within the sample [28]. NetMHCpan-4.1 was used to predict the binding affinity of peptides (8-14 amino acids in length) to select HLA class I molecules. These peptides were extracted from the immunopeptidome data and submitted to the tool using default parameters. NetMHCpan-4.1 employs a ranking system where a predicted rank ≤2 and >0.5 indicates a weak binder, while a score ≤0.5 indicates a strong binder [29].

### Validation of Predicted HLA Binders Using External Cancer and Normal Proteomes

Predicted HLA binders (rank ≤2) were searched against external cancer and normal tissue proteomics data using the web interface of PepQuery [30]. The search spanned 22 cancer proteomics cohorts (2,704 samples) and two normal tissue proteomics datasets (PXD010154 and PXD016999, about 30 tissue types, 242 samples). GENe annotation COmpilation and DEscription (GENCODE) version 34 Human was set as the reference background for the search. PepQuery hits where the provided peptides scored higher than any reference background and random peptide were selected (filtering for n_db = 0 and n_random = 0).

### Genomic Origins of *De Novo* Peptides

To determine the genomic origin of *de novo* peptides, we first translated the human reference genome and transcriptome (GRCh38.p14, GCF_000001405.40) across all six reading frames and combined the results with the human reference proteome to generate all possible translated peptides. *De novo* peptides were then searched against these databases using BLASTP [31] variant optimized for short peptides (blastp-short). Hits with at most one gap or one mismatch in the alignment (when considering the entire length of the *de novo* peptides) were selected. A *de novo* peptide was classified as originating from a known protein or transcript if there was a hit against the reference proteome or translated transcriptome. *De novo* peptides with no hits against the reference proteome and translated transcriptome but with hits against the translated genome were interpreted based on the functional annotations of their genomic coordinates. *De novo* peptides with no hits across all databases were classified as noncanonical peptides, potentially arising from cryptic ORFs, aberrant translation, RNA editing, or tumor-specific genetic alterations.

### Validation Using Synthetic Peptides

Synthetic peptides for spectra validation were ordered from GL Biochem (Shanghai) Ltd., as HPLC grade (≥50% purity). These were analyzed using the same LC–MS/MS system and acquisition parameters as indicated above for the endogenous peptides.

### ELISPOT Assay

Short (9-mer) and long (25-mer) peptides, corresponding to both wild-type and mutant sequences, were synthesized by GenScript at HPLC grade with ≥90% purity. Peripheral blood mononuclear cells (PBMCs) were isolated from donors predicted to present tumor antigens (based on the preceding analyses), using a standard Ficoll density gradient centrifugation. The protocol was adapted from Cimen Bozkus et al., 2021 [32]. On Day 0, T cell culture was initiated using unfractionated PBMCs. The cells were stimulated with a cytokine cocktail containing GM-CSF, IL-4, and Flt3-L for 24 hours to promote the differentiation of distinct dendritic cell (DC) subsets. On Day 1, 100 µL of supernatant was removed from each well, and replaced with 100 µL of a 2× adjuvant solution containing peptides. The final concentrations per well were as follows: 10 µM R848, 100 ng/mL LPS, 10 ng/mL IL-1β, and 10 µg each of wild-type and mutant peptides (both 9-mer and 25-mer). Cells were incubated for 24 hours. For T cell expansion, every three days over a 10-day period, 100 µL of supernatant was removed from each well and replaced with 100 µL of a 2× cytokine feeding solution, yielding final concentrations of 10 IU/mL IL-2, 10 ng/mL IL-7, and 10 ng/mL IL-15. For the ELISPOT assay, 1 × 10⁵ expanded T cells were seeded per well onto pre-coated ELISPOT plates in complete medium (AIM-V supplemented with 2% human serum and 1% penicillin-streptomycin) and rested for 24 hours. Wild-type and mutant peptides were then added to the respective wells, and the plates were incubated for 48 hours. The resulting spots were developed and scanned using a C.T.L. ELISPOT reader and quantified using ImmunoSpot 5.0 software (Cellular Technology Limited).

### MHC Peptide Binding Assay

Candidate peptides listed in immunopeptidome identification from the *de novo* sequencing strategy result section (data is available in the supplementary table 4) in were assembled with HLA-A*11:01 and analyzed using the ProImmune REVEAL® MHC-Peptide Binding Assay to determine their level of incorporation into MHC molecules. Binding to MHC molecules was compared to that of a known T cell epitope, a positive control peptide with strong binding properties (30).

## RESULTS

### Patient Demographics and HLA Allele Characterization

All participants had a mean age of 62.8 years (range: 42-77 years), and tumor tissues in this study were classified into subtypes based on tissue pathology, as shown in Table 1. Specifically, there were 10 patients with clear cell renal cell carcinoma (ccRCC), 2 patients with papillary RCC (pRCC), and 1 patient with clear cell papillary RCC (ccpRCC). Table 1 also details other patient characteristics, including gender, ISUP grade, and tumor stage. Four-digit HLA alleles for HLA-A, HLA-B, and HLA-C were identified in all 13 patients by whole-exome sequencing (WES) of peripheral blood mononuclear cells (PBMCs), followed by analysis using the well-established computational tool OptiType. The most common HLA alleles identified were HLA-A*11:01 (69%), HLA-B*40:01 (30.8%), and HLA-C*07:02 (30.7%). When comparing the list of HLA alleles from this study with the reported HLA alleles in four comprehensive HLA ligand databases [33, 34], i.e., the Immune Epitope Database (IEDB), HLA Ligand Atlas, Immune Epitope (IE) Atlas, and SysteMHC Atlas, the HLA typing from these 13 patients showed an overlap of 37 distinct HLA class I alleles. Interestingly, we identified 7 HLA alleles that, to our knowledge, have not been previously reported in immunopeptidome datasets: HLA-A*11:29, HLA-A*42:02, HLA-B*27:04, HLA-B*15:12, HLA-B*15:75, HLA-B*38:02, and HLA-B*55:02.

**Table 1.** Patient Demographics and HLA Allele Characterization.

| Patients ID | Gender | Age | Histological Subtype | ISUP Grade | Tumor Stage | HLA typing |  |  |
| --- | --- | --- | --- | --- | --- | --- | --- | --- |
|  |  |  |  |  |  | A | B | C |
| RCC3 | M | 59 | ccRCC | 3 | I | *02:07 | *46:01 | *01:02 |
|  |  |  |  |  |  | *74:01 | *46:01 | *07:02 |
| RCC5 | M | 66 | ccpRCC | 1 | I | *11:01 | *40:01 | *03:04 |
|  |  |  |  |  |  | *24:02 | *13:01 | *07:02 |
| RCC6 | F | 52 | ccRCC | 2 | I | *11:01 | *40:01 | *07:02 |
|  |  |  |  |  |  | *02:01 | *51:01 | *14:02 |
| RCC8 | F | 69 | ccRCC | 1 | I | *11:01 | *40:01 | *03:02 |
|  |  |  |  |  |  | *31:01 | *27:04 | *03:04 |
| RCC9 | F | 53 | ccRCC | 1 | III | *01:01 | *15:12 | *03:03 |
|  |  |  |  |  |  | *11:01 | *57:01 | *03:04 |
| RCC10 | F | 76 | pRCC | 2 | I | *11:01 | *15:32 | *03:02 |
|  |  |  |  |  |  | *33:03 | *58:01 | *12:03 |
| RCC11 | F | 71 | ccRCC | 2 | I | *02:07 | *07:05 | *15:01 |
|  |  |  |  |  |  | *29:01 | *51:01 | *15:02 |
| RCC12 | M | 72 | pRCC | 3 | I | *02:01 | *15:01 | *07:02 |
|  |  |  |  |  |  | *33:03 | *15:75 | *12:02 |
| RCC13 | M | 65 | ccRCC | 1 | III | *02:01 | *40:01 | *03:04 |
|  |  |  |  |  |  | *11:01 | *51:01 | *04:02 |
| RCC15 | M | 42 | ccRCC | 2 | III | *11:01 | *51:01 | *04:03 |
|  |  |  |  |  |  | *24:07 | *55:02 | *14:02 |
| RCC17 | F | 52 | ccRCC | 4 | I | *02:01 | *15:02 | *08:01 |
|  |  |  |  |  |  | *11:01 | *15:18 | *08:01 |
| RCC18 | F | 63 | ccRCC | 3 | III | *02:03 | *38:02 | *01:02 |
|  |  |  |  |  |  | *24:02 | *46:01 | *07:02 |
| RCC19 | M | 77 | ccRCC | 4 | IV | *11:02 | *13:02 | *06:02 |
|  |  |  |  |  |  | *11:29 | *27:04 | *12:02 |

### Analysis of Personalized Immunopeptidomes

Fig. 1 outlines the simplified workflow for personalized immunopeptidome profiling. Peripheral blood mononuclear cells (PBMCs) and tumor tissues were collected from each patient. Whole-exome sequencing (WES) was conducted on both PBMC and tumor samples, generating paired sequencing data. Germline variants from PBMCs and somatic variants from tumor samples were identified and used to construct a personalized proteome database for each patient by integrating these variants with reference proteome sequences from the UniProt Homo sapiens database. Tumor tissues underwent HLA class I immunoprecipitation to isolate HLA-bound peptides, which were analyzed via liquid chromatography-tandem mass spectrometry (LC-MS/MS). The resulting LC-MS/MS data were processed using two proteomics analysis programs: PEAKS Xpro and SMSNet. Each program performed two analyses for peptide identification: (1) a database search using the personalized proteome database and (2) *de novo* sequencing.

**Fig. 1.**
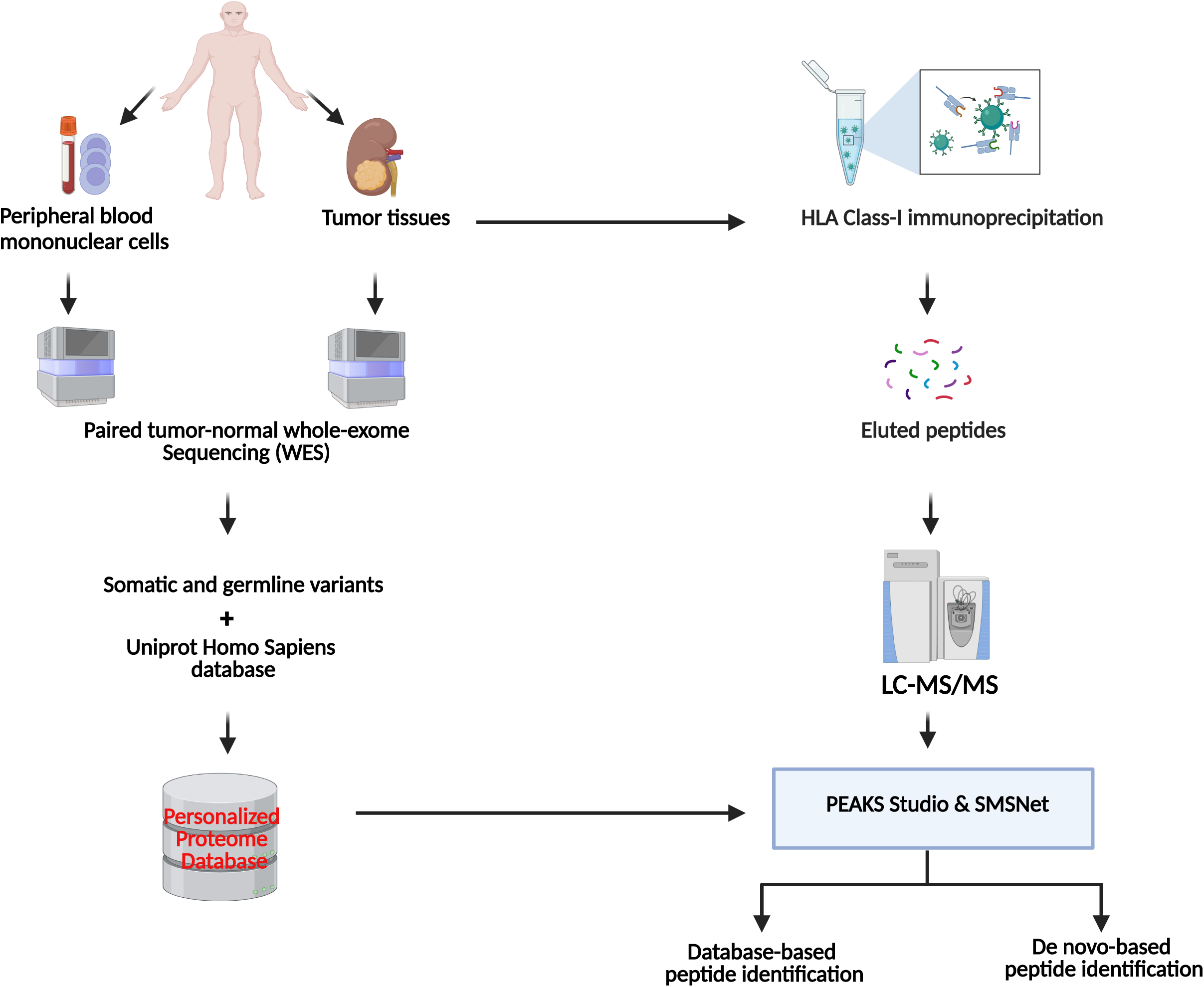
The workflow for personalized immunopeptidome profiling.

After constructing the personalized proteome database, we analyzed the types and frequencies of mutations in each patient. Whole-exome sequencing (WES) revealed that the majority of single nucleotide variants (SNVs) were missense mutations, accounting for 64.3%. Synonymous SNVs were the second most common, representing 23.1%. In addition to SNVs, we identified small insertions and deletions (indels), including in-frame deletions (3%) and in-frame insertions (0.1%), as well as stop-gain mutations (3%), as shown in Fig. 2a. These diverse mutation classes contributed to distinct mutational profiles across patients, as illustrated in Fig. 2b (the dataset for all patients is available in Table S1). Using WES data, we calculated the tumor mutational burden (TMB) for each patient, as shown in Fig. 2c. The TMB varied significantly, with RCC5 exhibiting the highest value.

**Fig. 2.**
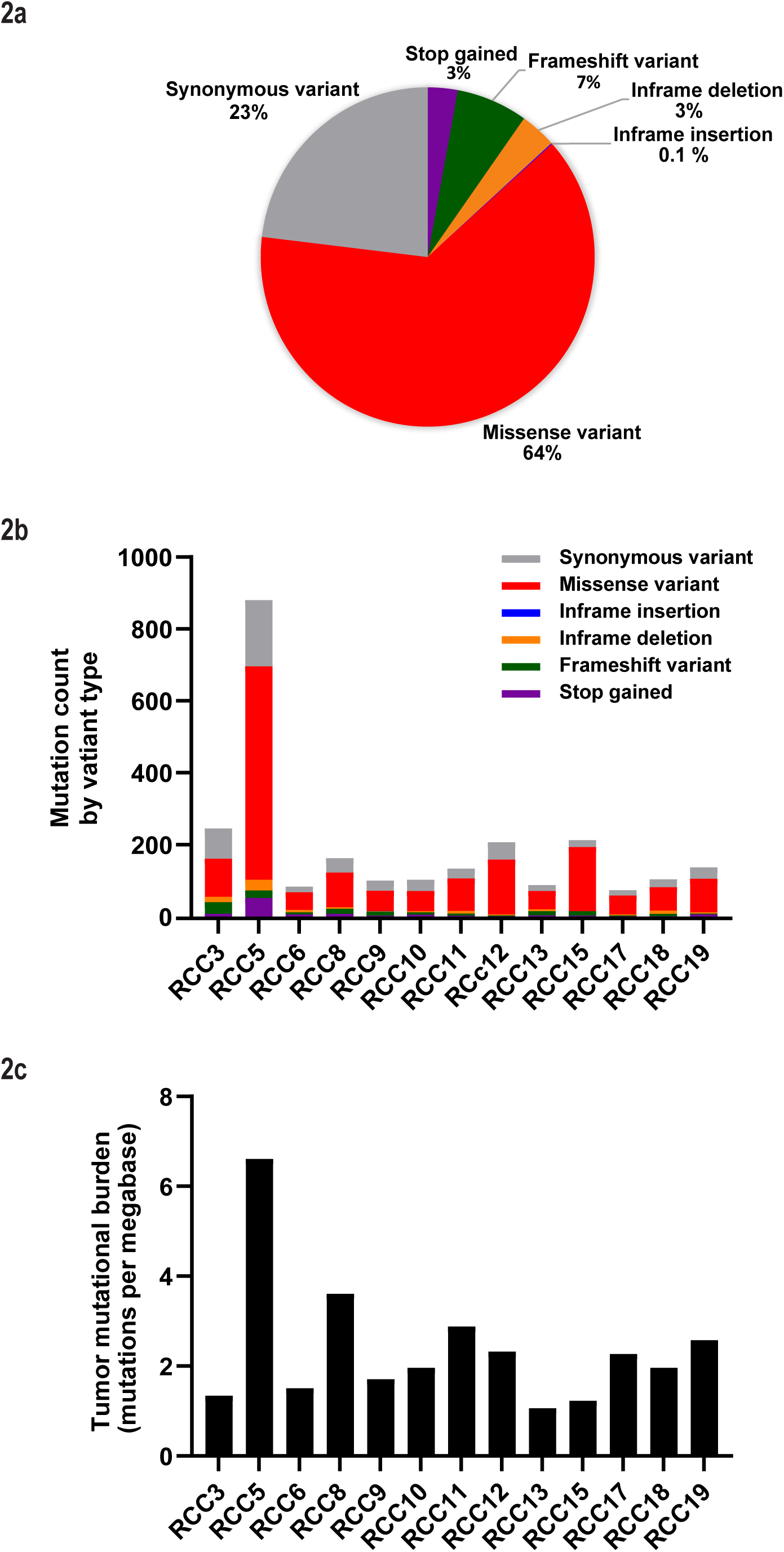
Mutation landscape and tumor mutational burden in RCC patients. a) Proportion of mutation types across RCC patients. The pie chart illustrates the distribution of mutations identified in the cohort. b) Mutation classifications by patients. The stacked bar chart shows the number and distribution of mutation types for each RCC patient. c) Tumor mutational burden (TMB) across RCC patients. The bar chart presents the TMB (Mutations per Megabase, Mut/Mb) for each patient.

### Immunopeptidome identification from the database search strategy using the personalized proteome database

Fig. 3a presents the combined results of our LC-MS/MS database search, integrating peptide identifications from both PEAKS Xpro and SMSNet. Across all patients, we identified between 8,978 and 55,862 peptides (the dataset for all patients is available in Table S2). As shown in Fig. 3b, the number of identified peptides is positively correlated with the amount of tissue analyzed (r = 0.51, P = 0.037), indicating that more tissue generally yields more HLA-bound peptides. Analysis of peptide length (Fig. 3c) reveals that most peptides range from 8 to 15 amino acids, with a predominant length of 9 amino acids—typical for HLA class I peptides. This confirms that our experimental approach effectively captures HLA class I–restricted peptides. We then used the Gibbs Clustering algorithm to identify characteristic peptide motifs for each patient, followed by matching these motifs to HLA allotype binding preferences using the NetMHC algorithm (see Methods for details on matching criteria). This analysis revealed that each patient exhibited 2–5 distinct peptide motifs corresponding to their HLA allotype binding preference clusters, demonstrating that in most patients, these motifs align with at least 50% of their HLA allotypes (Fig. 3d). Notably, some expected binding motifs were missing for certain alleles. A likely explanation is that an insufficient number of peptides was identified for those alleles, which is supported by the positive correlation between the number of matched motifs and the overall number of peptides detected per patient. Fig. 3e provides a representative example from patient RCC13, illustrating the observed motifs and their corresponding HLA allotype binding preference clusters. The complete dataset for all patients is available in Fig. S1.

**Fig. 3.**
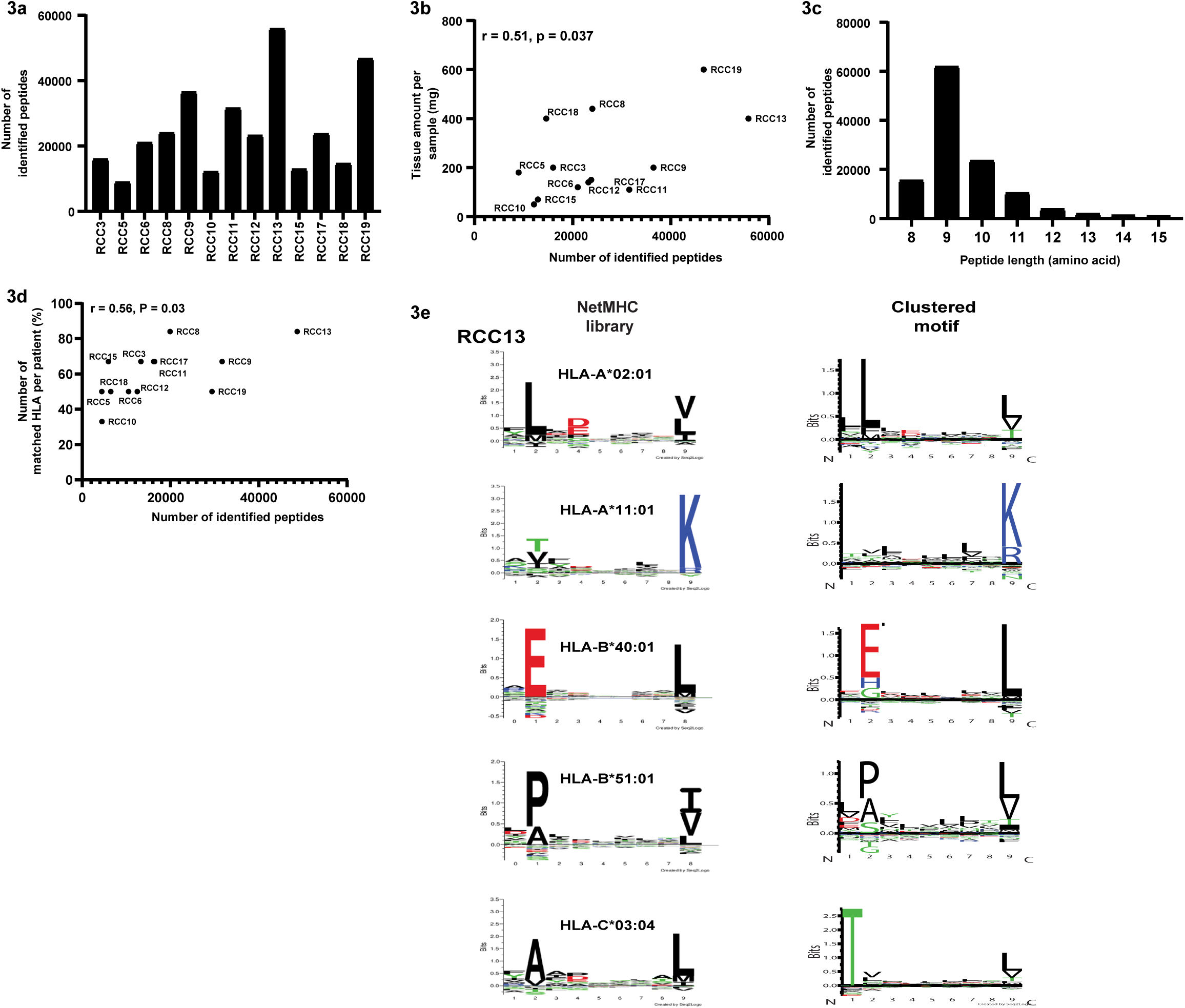
Peptide identification, distribution, and HLA binding motifs in RCC patients. a) The bar chart shows the total number of peptides identified in each RCC patient sample through mass spectrometry analysis. b) A scatterplot showing the correlation between the amount of tissue used per sample (mg) and the number of identified peptides. c) A histogram depicting the length distribution of identified peptides. d) A scatterplot illustrating a correlation between the percentage of matched HLA alleles and the number of identified peptides per patient. e) Representative clustered motifs and HLA allotype binding preferences in RCC13. Motifs generated using GibbsCluster ("Clustered Motif") are shown alongside NetMHC library predictions for HLA allotypes in patient RCC13. Each panel represents a distinct HLA allele, with sequence logos indicating amino acid binding preferences for each motif. Larger letters in the sequence logos represent higher conservation or preference at a given position. The complete dataset, including motifs for all patients, is available in Supplementary Fig. 1.

### Identification of a Neoantigen and Its Immune Response Using a Database Search Strategy

Among the 13 RCC patients analyzed for their immunopeptidome using mass spectrometry, a single neoantigen was identified. Our results revealed a mutant JADE2–derived peptide corresponding to a threonine-to-serine substitution at protein position 391 in the RCC5 patient, who exhibited the highest tumor mutational burden (TMB). Computational analysis using NetMHCpan-4.1 predicted strong binding of the mutant peptide to HLA-B*40:01 (Fig. 4a). To validate this prediction, we synthesized the mutant peptide and performed mass spectrometry analysis. Comparison of LC-MS/MS spectra and retention time (RT) between the synthetic peptide and the endogenous mutant peptide identified by immunopeptidomics confirmed successful identification of the mutant peptide (Fig. 4b). To evaluate the immunogenicity of the identified mutant peptide and assess mutation-specific T-cell responses in RCC5, we conducted an ELISPOT assay using PBMCs from the RCC5 patient and a healthy donor matched for HLA-B*40:01. PBMCs were co-cultured with both short (9 amino acids) and long (25 amino acids) versions of the mutant and wild-type peptides, and IFN-γ production was measured. The ELISPOT assay revealed a significantly higher number of IFN-γ-producing T cells in response to the mutant peptide compared to the wild-type peptide in the RCC5 patient’s PBMCs. Importantly, neither the mutant nor wild-type peptide elicited a response in the healthy donor’s PBMCs (Fig. 4c). These findings suggest that the mutant JADE2 peptide can specifically stimulate T cells from the RCC5 patient, underscoring its potential as a tumor-specific neoantigen candidate. The lack of response in the normal control PBMCs further supports the patient-specific nature of this immune recognition, highlighting its relevance for patient-specific immunotherapeutic strategies.

**Fig. 4.**
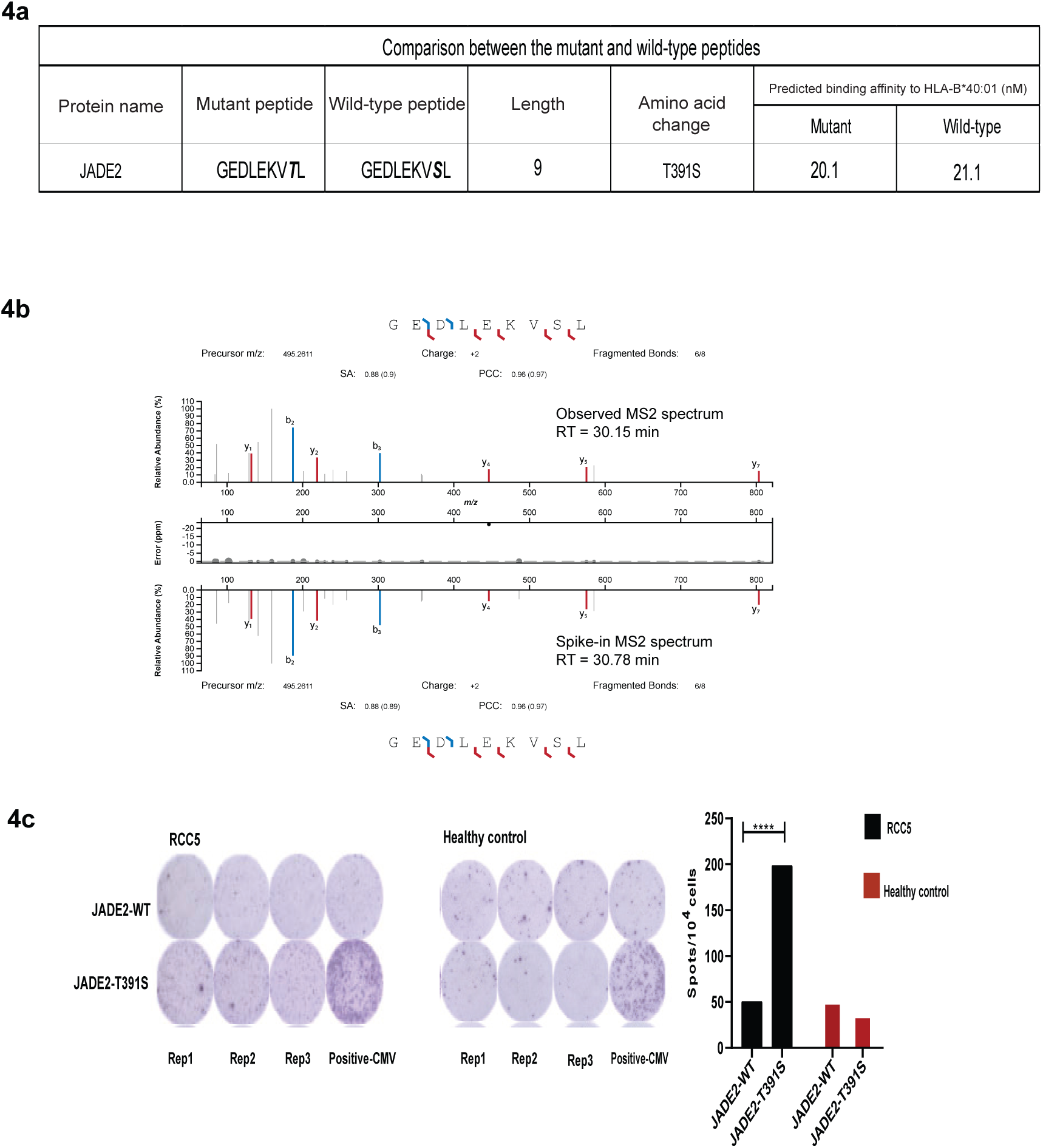
Identification and validation of a JADE2 neoantigen. a) A T391S substitution in JADE2 generates the mutant-derived peptide GEDLKEVTL with predicted strong binding to HLA-B*40:01. b) The endogenous mutant peptide was confirmed by matching MS2 spectra and retention time with the synthetic peptide. c) IFN-γ ELISPOT showed a strong mutant-specific T-cell response in PBMCs from the RCC5 patient, with no response observed in an HLA-matched healthy donor.

### Immunopeptidome Identification from the *De Novo* Sequencing Strategy

To overcome the limitations of the database search strategy, which is restricted to peptides present in the personalized proteome database, we employed a *de novo* sequencing approach to identify tumor peptides derived from non-canonical mechanisms, such as splice variants, intron retention, novel ORFs, and retroelements which may not be captured by whole-exome sequencing [35]. Using this approach, we identified between 6,693 and 20,863 peptides per RCC patient sample that were not present in the personalized proteome database (dataset for all patients is available in Table S3) with a positive correlation observed between the number of identified peptides and the amount of tissue analyzed (Fig. 5a and 5b). The majority of peptides identified were between 8 and 15 amino acids in length, with a predominant length of 9 amino acids, consistent with MHC class I binding preferences (Fig. 5c). Motifs generated using Gibbs Clustering revealed 2–5 distinct motifs per patient, corresponding to their HLA allotype binding preferences, and these motifs aligned with at least 50% of the HLA allotypes in most patients (Fig. 5d). To validate these findings, 56 peptides classified in the HLA-A*11:01 cluster were randomly selected, synthesized, and analyzed using independent spike-in LC-MS/MS experiments (dataset is available in Table S4), showing strong agreement between 51 synthetic MS/MS spectra and their corresponding endogenous spectra and retention times (Fig. 5E); only five synthetic MS/MS spectra were not detected (Fig. S2). The binding affinity of these 51 peptides was then evaluated using the ProImmune REVEAL® MHC-peptide binding assay and compared with NetMHCpan-4.1 predictions, which identified 45 peptides as strong binders and 6 as weak binders as shown in pink (Fig. 5f). The ProImmune assay confirmed that 45/51 strong binders passed the assay threshold while 43/45 "passed" epitopes were also classified as strong binders by NetMHCpan-4.1 predictions. Interestingly, 2 of the 6 "non-passed" epitopes were also identified as strong binders by NetMHCpan-4.1. A significant negative correlation was observed between the NetMHCpan-4.1 binding affinity predictions and ProImmune REVEAL® assay scores (r = -0.26, P = 0.029). The *de novo* sequencing strategy enabled the identification of potential non-canonical tumor-derived peptides, validated through LC-MS/MS, and revealed a significant relationship between predicted and experimental binding affinities, providing strong evidence that these peptides are bona fide HLA binders. This highlights the utility of *de novo* sequencing in identifying novel epitopes beyond conventional database-based methods.

**Fig. 5.**
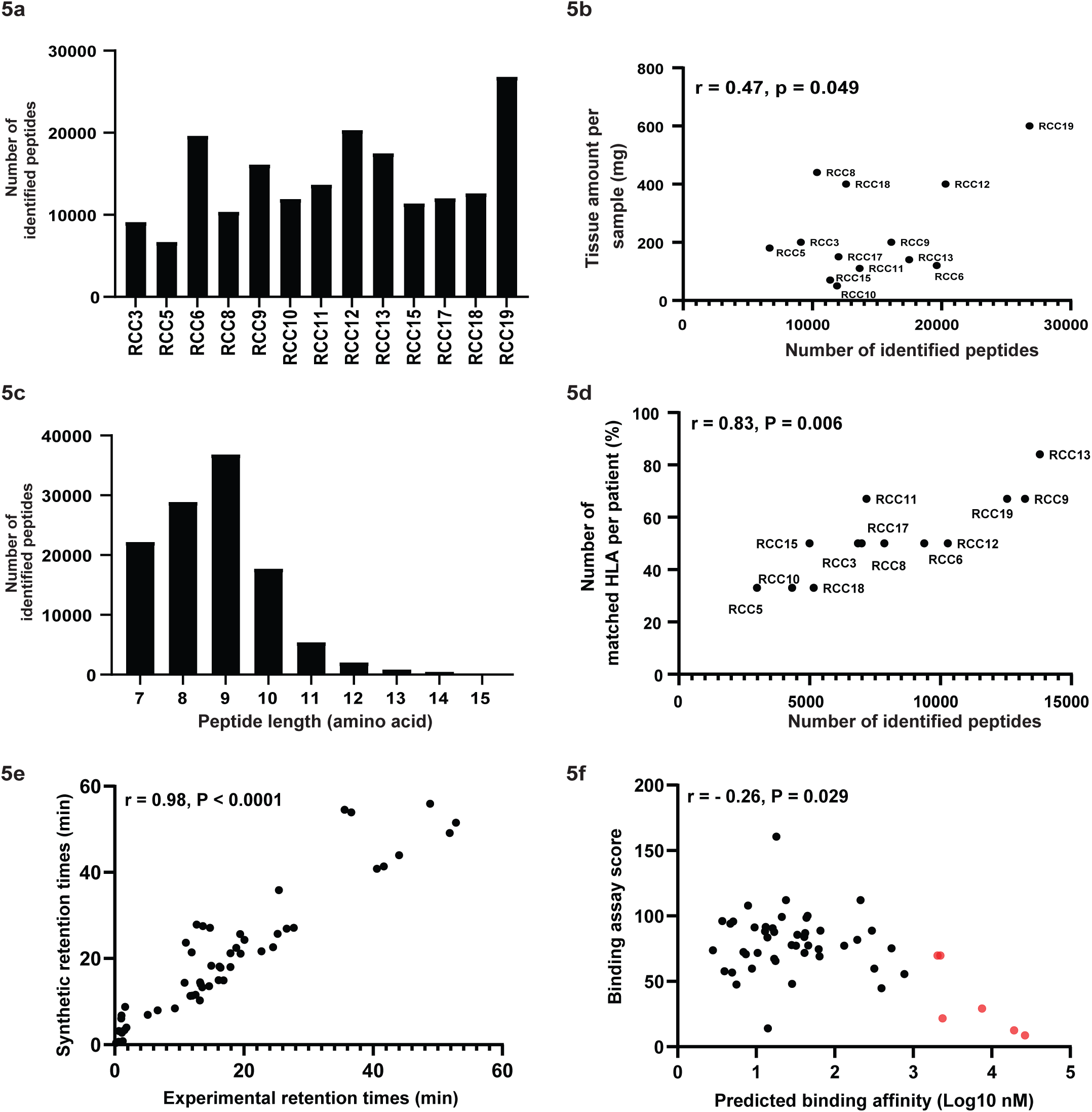
Immunopeptidome identification and validation using a *de novo* sequencing strategy. a) Number of *de novo*–identified peptides across 13 RCC samples. b) Positive correlation between tissue amount and number of identified peptides. c) Length distribution of *de novo*–identified peptides, with a predominant peak at 9 amino acids. d) Correlation between the proportion of HLA allotypes matched by GibbsCluster-derived motifs and the number of identified peptides. e) Validation of *de novo*–identified peptides by comparison of synthetic and experimental MS/MS spectra and retention times. f) Comparison between NetMHCpan-4.1 predicted binding affinities and ProImmune REVEAL® binding scores for 51 validated peptides, showing a significant negative correlation.

### Investigation of *De Novo*–Identified Tumor-Associated Antigen Candidates

To identify clinically relevant tumor-associated antigen candidates, we first selected 799 unique peptides derived from *de novo* sequencing that were detected in at least three patients. Subsequently, we performed MHC binding prediction for all patient-specific HLA-A and HLA-B alleles using NetMHCpan 4.1. Based on percentile rank thresholds (<0.5% for strong binders and <2% for weak binders), a total of 320 peptides were predicted to bind at least one HLA class I allele (Table S5). To further increase confidence in the predicted binders, we selected a subset of 241 peptides predicted to bind to at least two HLA alleles. Although this additional filtering step is optional, it enriches peptides with broader immunogenic potential. To evaluate immunopeptidome evidence supporting their tumor association, we searched for these peptides against large-scale cancer and normal proteomics datasets (Search Date: July 2025, see materials and methods) using PepQuery [30]. Among 241 candidate peptides, 35 were confidently detected in at least 3 cancer samples. Notably, the peptides SLMHAYLLK (found in 22 cancer samples), TALLDLLTK (16 cancer samples), and VENFEVDSL (13 cancer samples) were not detectable in normal tissue datasets, suggesting that these peptides may represent promising cancer-associated HLA-presented antigens.

Next, we further investigated the genomic origins of these 35 high-confidence *de novo*–identified cancer-associated peptides by mapping them against the human reference proteome, six-frame translations of the human reference genome, and the translated human reference transcriptome. Only perfect matches and near-perfect matches (at most one gap/mismatch) were considered to minimize false positives. This analysis aimed to distinguish peptides arising from canonical coding regions from those potentially derived from noncanonical ORFs, alternative splicing, or tumor-specific aberrant translations. There was only one perfect hit between peptide SVAWPFSFPK and VPS36 transcript isoforms (Table 2, NM_016075.3, NM_001282169.1, and NM_001282168.1) within a 5′ overlapping upstream open reading frame (uORF). There were five additional near-perfect hits to three known proteins and two non-coding regions. The lack of hits suggests that these *de novo*–identified peptides may originate from complex noncanonical events and potentially tumor-associated genomic alterations.

**Table 2.**
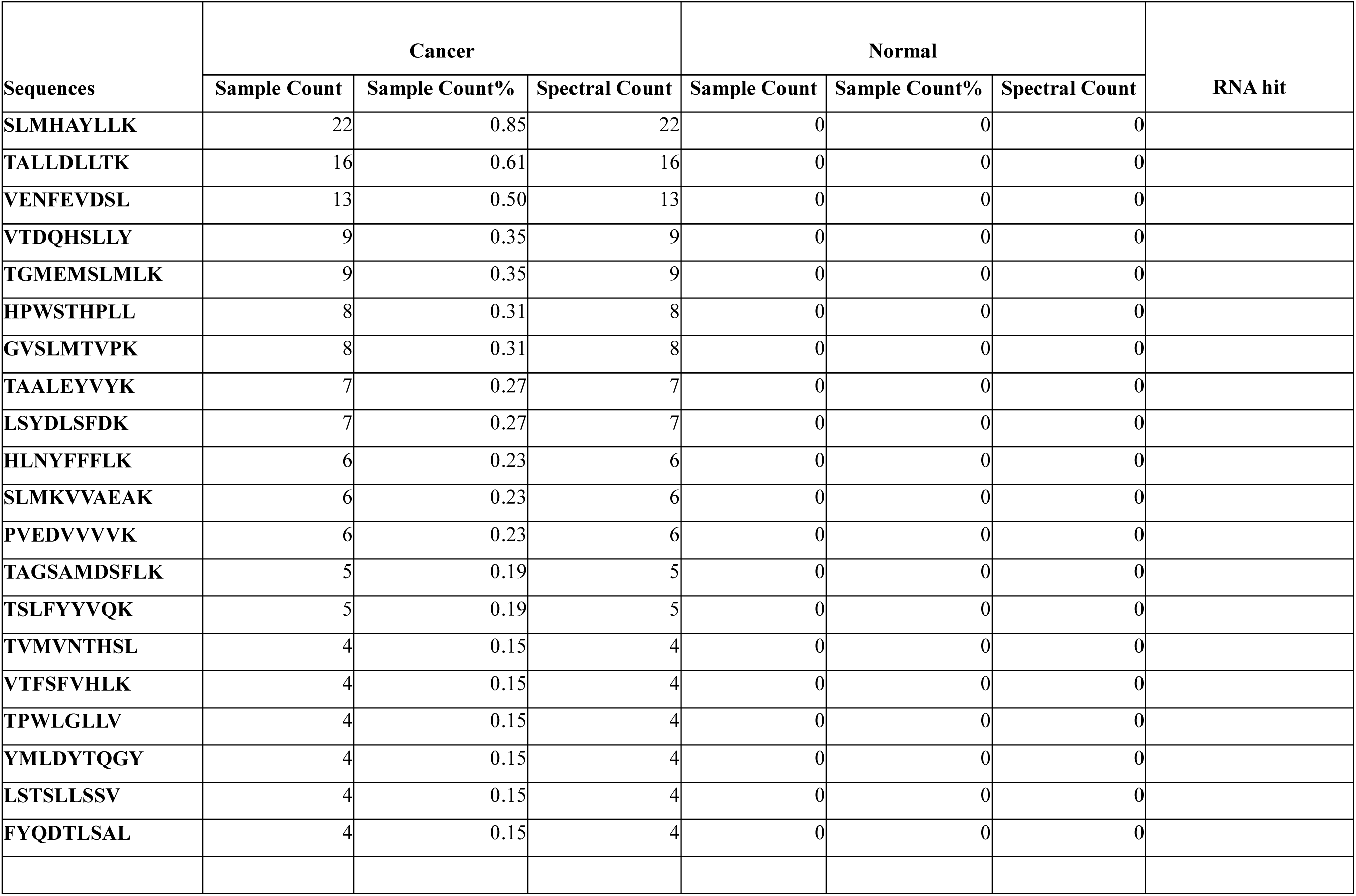
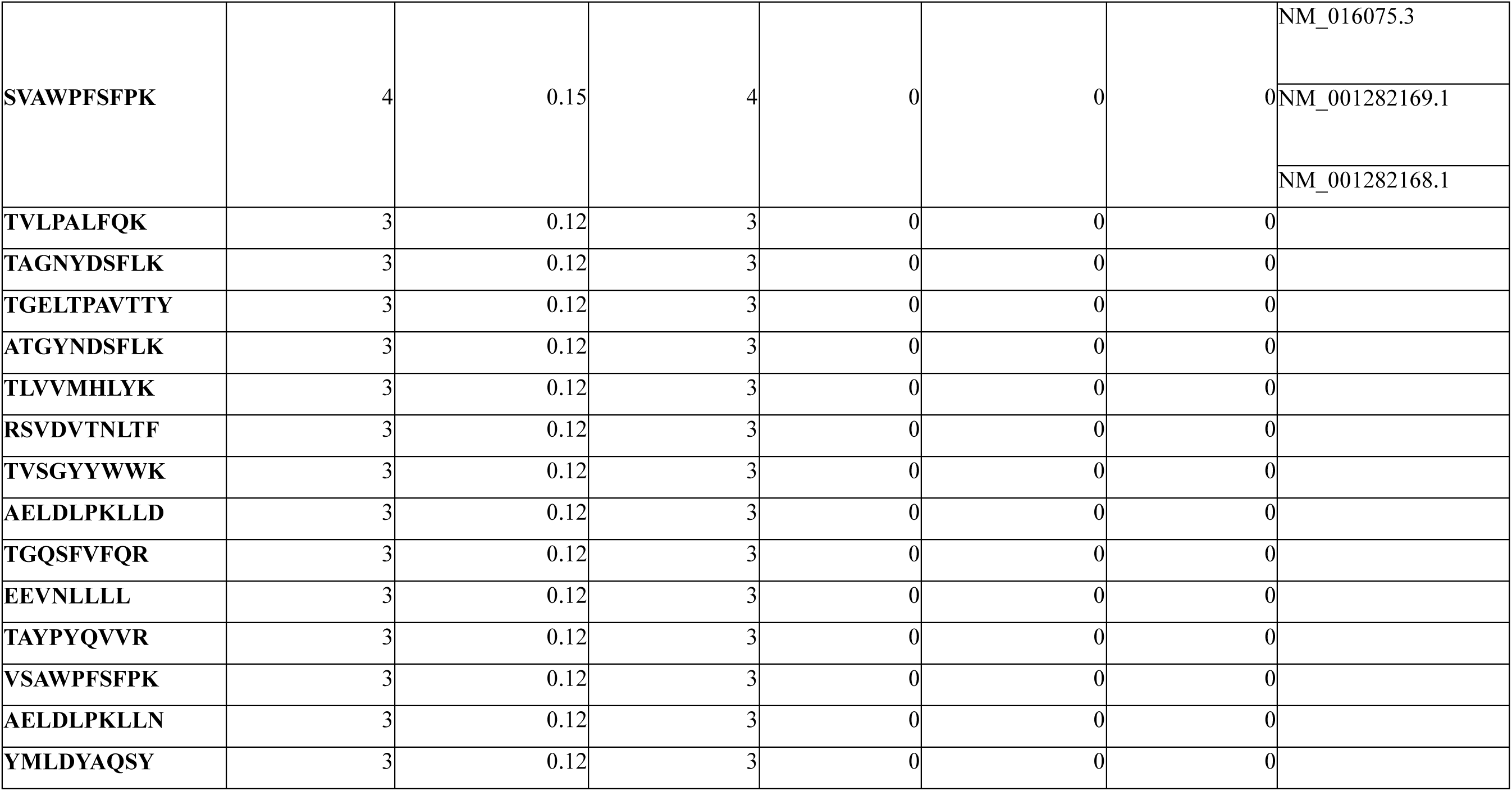
*De novo*–identified peptides supported by cancer proteomes and their genomic annotation.

## Discussion

This study advances our understanding of the personalized immunopeptidomes of Thai patients with renal cell carcinoma (RCC), a malignancy with substantial global impact. The research addresses a critical gap in the existing literature by focusing on a Southeast Asian population, which has been historically underrepresented in immunopeptidomic studies. By employing advanced mass spectrometry techniques and a combined strategy of database searching and *de novo* sequencing, the study provides a comprehensive characterization of HLA class I-presented peptides in this specific ethnic group, paving the way for the development of tailored immunotherapeutic approaches [36]. A finding of this research is the identification of seven HLA alleles that, to our knowledge, have not been previously reported in immunopeptidome datasets [37]. Because HLA alleles directly determine which peptides are presented to the immune system, population-specific HLA diversity is expected to shape the repertoire of tumor-associated antigens observed in different cohorts. In this study, we examined the distribution of common HLA class I alleles, including HLA-A*11:01, HLA-B*40:01, and HLA-C*07:02, within the Thai RCC cohort. Consistent with prior reports, HLA-A*11:01 was the most prevalent allele, reflecting its high frequency in East Asian populations, while HLA-B*40:01 and HLA-C*07:02 showed more variable representation across ethnic groups [38-41]. We also observed a positive correlation between the amount of tumor tissue analyzed and the number of HLA-bound peptides identified [10, 42], indicating that sample input remains a key determinant of immunopeptidome depth. This finding underscores the importance of sufficient starting material in immunopeptidomic studies to achieve a comprehensive view of the presented peptides. It also highlights the ongoing need for advancements in peptide extraction and enrichment methodologies, particularly when working with limited clinical samples.

The database search strategy led to the identification of a single neoantigen derived from the JADE2 gene in the RCC5 patient, who presented with the highest tumor mutational burden (TMB) in the cohort. This finding aligns with the understanding that higher TMB often correlates with an increased likelihood of generating neoantigens [43-45]. The JADE2 gene, involved in chromatin remodeling and cell cycle regulation, has been implicated in various cancers [46, 47]. The identification of a specific mutation in JADE2 leading to a neoantigen suggests its potential role in the tumor’s immunogenicity in this patient. However, the fact that only one neoantigen was identified in the highest TMB patient indicates that other factors beyond mutation load, such as the nature and location of mutations and the HLA binding affinity of the resulting peptides, are critical determinants of neoantigen presentation.

The identification of tumor-specific neoantigens and tumor-associated antigens is central to the development of personalized cancer immunotherapies tailored to individual tumor antigen landscapes and HLA genotypes [10, 11, 48]. In this study, we applied a *de novo* sequencing–based approach to uncover novel peptide sequences that may not be represented in conventional protein databases [49]. We further validated the *de novo*–identified peptides, specifically, 56 candidates from the HLA-A*11:01 cluster were synthesized and analyzed by independent spike-in LC–MS/MS experiments. High concordance between experimental and synthetic MS/MS spectra and retention times was observed for 51 peptides, supporting the accuracy of *de novo* sequencing when combined with stringent spectral validation, while the absence of five peptides likely reflects stochastic sampling or reduced ionization efficiency inherent to shotgun MS analyses. Functional relevance was further supported by MHC binding assays, in which 45 of 51 peptides predicted as strong binders by NetMHCpan-4.1 were confirmed by the ProImmune REVEAL® assay. A significant negative correlation between predicted binding affinity and experimental binding score was observed, consistent with previous reports indicating that discrepancies between computational predictions and experimental binding can arise from factors such as peptide processing, HLA stability, and assay-specific sensitivity. The high concordance between synthetic and experimental MS/MS spectra for the majority of *de novo*–identified peptides underscores the robustness of *de novo* sequencing when supported by stringent spectral validation. Importantly, the inability to detect a small subset of peptides is consistent with known technical limitations of shotgun proteomics, such as stochastic precursor selection and variable ionization efficiency, rather than inaccuracies in sequence inference. Furthermore, the strong agreement between NetMHCpan-4.1 predictions and ProImmune REVEAL® binding results supports the biological relevance of these peptides as bona fide HLA ligands. The modest but significant inverse correlation between predicted binding affinity and experimental binding scores highlights the complementary nature of computational and experimental approaches, as discrepancies likely reflect additional layers of antigen processing and presentation that are not fully captured by in silico models alone. Collectively, these findings reinforce the value of integrating *de novo* sequencing with orthogonal validation strategies to confidently identify functional HLA-presented peptides. Our approach aligns with recent studies that advocate the integration of *de novo* sequencing with spectral validation tools to overcome the limitations of reference-based proteomic searches. For example, Wen et al. (2023) demonstrated the utility of PepQuery2 in large-scale validation of non-canonical peptides using over a billion indexed spectra from cancer datasets. Similarly, Liao et al. (2024) applied *de novo*–assisted discovery to identify MHC-bound peptides derived from noncoding or unannotated regions in cancer.

While traditional proteogenomic pipelines typically rely on RNA-seq or whole-exome sequencing for database construction, our method bypasses the need for transcriptomic data and allows for direct capture of the immunopeptidome at the protein level. This is particularly valuable in clinical settings where tumor material is limited or RNA is degraded. In conclusion, the combination of *de novo* sequencing and PepQuery validation offers a robust framework for identifying tumor-associated peptides, particularly in settings where conventional methods fall short. This approach holds promise for broadening the scope of neoantigen and tumor-associated antigen discovery, especially in underrepresented populations and cancers with low mutational burden. A significant strength of this study lies in the successful implementation of *de novo* sequencing, which allowed for the identification of a large number of peptides not found in the personalized proteome database. This approach is crucial for discovering non-canonical tumor antigens arising from events that may not be captured by whole-exome sequencing, such as splice variants or novel open reading frames. The rigorous validation of these *de novo*-identified peptides through synthetic peptide experiments and HLA binding assays provides strong evidence for their authenticity and their potential to be presented by HLA molecules, significantly expanding the scope of potential tumor antigens. However, current immunopeptidomic approaches face significant challenges, including low detection sensitivity, stochastic peptide sampling, difficulty in detecting rare or low-abundance peptides, and the need for robust orthogonal validation to confirm tumor association. Overcoming these methodological barriers will be essential for systematically uncovering cancer-specific peptides generated from non-canonical translation.

## Acknowledgments

This study was supported by the Thailand Science Research and Innovation Fund, Chulalongkorn University (HEA663000034, HEA663000041 and HEAF673000102) and National Research Council of Thailand (NRCT), and Chulalongkorn University (N42A670716). The academic endeavors of the Thailand Hub of Talents in Cancer Immunotherapy (TTCI) are supported by the National Research Council of Thailand. We sincerely thank all members of the Center of Excellence in Systems Biology for their valuable comments and constructive advice. We also gratefully acknowledge Boyang Zhao for his technical support in neoantigen calling.

## Contributions

Conception and design of study: T.P. and P.S.; Sample collection: T.A., J.O., S.S. and S.N.; Sample preparation: S.N., P.M. and P.S.; Acquisition of data: P.S., P.W., S.S., S.M., T.K. and P.C.; Analysis and/or interpretation of data: P.S., S.S., J.T., P.K. and T.P.; Drafting the manuscript: P.S., S.S., P.K., J.T., P.M., S.N., T.A., J.O., N.H., S.M., T.K., P.C. and T.P.; Approval of the version of the manuscript to be published: P.S., S.S., P.K., J.T., P.M., S.N., T.A., J.O., S.S., N.H. and T.P.

## Ethics declarations

The authors declare no competing interests.

## References

1. Cai Q, Chen Y, Qi X, Zhang D, Pan J, Xie Z, et al. Temporal trends of kidney cancer incidence and mortality from 1990 to 2016 and projections to 2030. Transl Androl Urol. 2020;9(2):166–81. doi: 10.21037/tau.2020.02.23. PubMed PMID: 32420123; PubMed Central PMCID: PMCPMC7215038.

2. Samnani S, Sachedina F, Gupta M, Guo E, Navani V. Mechanisms and clinical implications in renal carcinoma resistance: narrative review of immune checkpoint inhibitors. Cancer Drug Resist. 2023;6(2):416–29. Epub 20230627. doi: 10.20517/cdr.2023.02. PubMed PMID: 37457122; PubMed Central PMCID: PMCPMC10344724.

3. Ott PA, Hu Z, Keskin DB, Shukla SA, Sun J, Bozym DJ, et al. An immunogenic personal neoantigen vaccine for patients with melanoma. Nature. 2017;547(7662):217–21. Epub 20170705. doi: 10.1038/nature22991. PubMed PMID: 28678778; PubMed Central PMCID: PMCPMC5577644.

4. Sahin U, Derhovanessian E, Miller M, Kloke BP, Simon P, Lower M, et al. Personalized RNA mutanome vaccines mobilize poly-specific therapeutic immunity against cancer. Nature. 2017;547(7662):222–6. Epub 20170705. doi: 10.1038/nature23003. PubMed PMID: 28678784.

5. Keskin DB, Anandappa AJ, Sun J, Tirosh I, Mathewson ND, Li S, et al. Neoantigen vaccine generates intratumoral T cell responses in phase Ib glioblastoma trial. Nature. 2019;565(7738):234–9. Epub 20181219. doi: 10.1038/s41586-018-0792-9. PubMed PMID: 30568305; PubMed Central PMCID: PMCPMC6546179.

6. Shapiro IE, Bassani-Sternberg M. The impact of immunopeptidomics: From basic research to clinical implementation. Semin Immunol. 2023;66:101727. Epub 20230208. doi: 10.1016/j.smim.2023.101727. PubMed PMID: 36764021.

7. Chen F, Zou Z, Du J, Su S, Shao J, Meng F, et al. Neoantigen identification strategies enable personalized immunotherapy in refractory solid tumors. J Clin Invest. 2019;129(5):2056–70. Epub 20190305. doi: 10.1172/JCI99538. PubMed PMID: 30835255; PubMed Central PMCID: PMCPMC6486339.

8. Chowell D, Krishna C, Pierini F, Makarov V, Rizvi NA, Kuo F, et al. Evolutionary divergence of HLA class I genotype impacts efficacy of cancer immunotherapy. Nat Med. 2019;25(11):1715–20. Epub 20191107. doi: 10.1038/s41591-019-0639-4. PubMed PMID: 31700181; PubMed Central PMCID: PMCPMC7938381.

9. Marcu A, Bichmann L, Kuchenbecker L, Kowalewski DJ, Freudenmann LK, Backert L, et al. HLA Ligand Atlas: a benign reference of HLA-presented peptides to improve T-cell-based cancer immunotherapy. J Immunother Cancer. 2021;9(4). doi: 10.1136/jitc-2020-002071. PubMed PMID: 33858848; PubMed Central PMCID: PMCPMC8054196.

10. Bassani-Sternberg M, Braunlein E, Klar R, Engleitner T, Sinitcyn P, Audehm S, et al. Direct identification of clinically relevant neoepitopes presented on native human melanoma tissue by mass spectrometry. Nat Commun. 2016;7:13404. Epub 20161121. doi: 10.1038/ncomms13404. PubMed PMID: 27869121; PubMed Central PMCID: PMCPMC5121339.

11. Shraibman B, Barnea E, Kadosh DM, Haimovich Y, Slobodin G, Rosner I, et al. Identification of Tumor Antigens Among the HLA Peptidomes of Glioblastoma Tumors and Plasma. Mol Cell Proteomics. 2019;18(6):1255–68. doi: 10.1074/mcp.RA119.001524. PubMed PMID: 31154438; PubMed Central PMCID: PMCPMC6553928.

12. Schuster H, Peper JK, Bosmuller HC, Rohle K, Backert L, Bilich T, et al. The immunopeptidomic landscape of ovarian carcinomas. Proc Natl Acad Sci U S A. 2017;114(46):E9942-E51. Epub 20171101. doi: 10.1073/pnas.1707658114. PubMed PMID: 29093164; PubMed Central PMCID: PMCPMC5699044.

13. Minegishi Y, Kiyotani K, Nemoto K, Inoue Y, Haga Y, Fujii R, et al. Differential ion mobility mass spectrometry in immunopeptidomics identifies neoantigens carrying colorectal cancer driver mutations. Commun Biol. 2022;5(1):831. Epub 20220818. doi: 10.1038/s42003-022-03807-w. PubMed PMID: 35982173; PubMed Central PMCID: PMCPMC9388627.

14. Arrieta-Bolanos E, Hernandez-Zaragoza DI, Barquera R. An HLA map of the world: A comparison of HLA frequencies in 200 worldwide populations reveals diverse patterns for class I and class II. Front Genet. 2023;14:866407. Epub 20230323. doi: 10.3389/fgene.2023.866407. PubMed PMID: 37035735; PubMed Central PMCID: PMCPMC10076764.

15. Sricharoensuk C, Boonchalermvichien T, Muanwien P, Somparn P, Pisitkun T, Sriswasdi S. Unsupervised Mining of HLA-I Peptidomes Reveals New Binding Motifs and Potential False Positives in the Community Database. Front Immunol. 2022;13:847756. Epub 20220321. doi: 10.3389/fimmu.2022.847756. PubMed PMID: 35386688; PubMed Central PMCID: PMCPMC8977642.

16. S A. FastQC: a quality control tool for high throughput sequence data 2010. Available from: https://www.bioinformatics.babraham.ac.uk/projects/fastqc/.

17. Li H, Durbin R. Fast and accurate short read alignment with Burrows-Wheeler transform. Bioinformatics. 2009;25(14):1754–60. Epub 20090518. doi: 10.1093/bioinformatics/btp324. PubMed PMID: 19451168; PubMed Central PMCID: PMCPMC2705234.

18. Benjamin D ST, Cibulskis K, Getz G, Stewart C, Lichtenstein L. Calling somatic SNVs and indels with Mutect2. bioRxiv. 2019;861054. doi: 10.1101/861054.

19. Koboldt DC, Zhang Q, Larson DE, Shen D, McLellan MD, Lin L, et al. VarScan 2: somatic mutation and copy number alteration discovery in cancer by exome sequencing. Genome Res. 2012;22(3):568–76. Epub 20120202. doi: 10.1101/gr.129684.111. PubMed PMID: 22300766; PubMed Central PMCID: PMCPMC3290792.

20. Kim S, Scheffler K, Halpern AL, Bekritsky MA, Noh E, Kallberg M, et al. Strelka2: fast and accurate calling of germline and somatic variants. Nat Methods. 2018;15(8):591–4. Epub 20180716. doi: 10.1038/s41592-018-0051-x. PubMed PMID: 30013048.

21. Szolek A, Schubert B, Mohr C, Sturm M, Feldhahn M, Kohlbacher O. OptiType: precision HLA typing from next-generation sequencing data. Bioinformatics. 2014;30(23):3310–6. Epub 20140820. doi: 10.1093/bioinformatics/btu548. PubMed PMID: 25143287; PubMed Central PMCID: PMCPMC4441069.

22. Dilthey AT, Mentzer AJ, Carapito R, Cutland C, Cereb N, Madhi SA, et al. HLA*LA-HLA typing from linearly projected graph alignments. Bioinformatics. 2019;35(21):4394–6. doi: 10.1093/bioinformatics/btz235. PubMed PMID: 30942877; PubMed Central PMCID: PMCPMC6821427.

23. Chalmers ZR, Connelly CF, Fabrizio D, Gay L, Ali SM, Ennis R, et al. Analysis of 100,000 human cancer genomes reveals the landscape of tumor mutational burden. Genome Med. 2017;9(1):34. Epub 20170419. doi: 10.1186/s13073-017-0424-2. PubMed PMID: 28420421; PubMed Central PMCID: PMCPMC5395719.

24. Purcell AW, Ramarathinam SH, Ternette N. Mass spectrometry-based identification of MHC-bound peptides for immunopeptidomics. Nat Protoc. 2019;14(6):1687–707. Epub 20190515. doi: 10.1038/s41596-019-0133-y. PubMed PMID: 31092913.

25. Perez-Riverol Y, Bai J, Bandla C, Garcia-Seisdedos D, Hewapathirana S, Kamatchinathan S, et al. The PRIDE database resources in 2022: a hub for mass spectrometry-based proteomics evidences. Nucleic Acids Res. 2022;50(D1):D543–D52. doi: 10.1093/nar/gkab1038. PubMed PMID: 34723319; PubMed Central PMCID: PMCPMC8728295.

26. Adusumilli R, Mallick P. Data Conversion with ProteoWizard msConvert. Methods Mol Biol. 2017;1550:339–68. doi: 10.1007/978-1-4939-6747-6_23. PubMed PMID: 28188540.

27. Karunratanakul K, Tang HY, Speicher DW, Chuangsuwanich E, Sriswasdi S. Uncovering Thousands of New Peptides with Sequence-Mask-Search Hybrid De Novo Peptide Sequencing Framework. Mol Cell Proteomics. 2019;18(12):2478–91. Epub 20191007. doi: 10.1074/mcp.TIR119.001656. PubMed PMID: 31591261; PubMed Central PMCID: PMCPMC6885704.

28. Andreatta M, Nielsen M. Gapped sequence alignment using artificial neural networks: application to the MHC class I system. Bioinformatics. 2016;32(4):511–7. Epub 20151029. doi: 10.1093/bioinformatics/btv639. PubMed PMID: 26515819; PubMed Central PMCID: PMCPMC6402319.

29. Reynisson B, Alvarez B, Paul S, Peters B, Nielsen M. NetMHCpan-4.1 and NetMHCIIpan-4.0: improved predictions of MHC antigen presentation by concurrent motif deconvolution and integration of MS MHC eluted ligand data. Nucleic Acids Res. 2020;48(W1):W449–W54. doi: 10.1093/nar/gkaa379. PubMed PMID: 32406916; PubMed Central PMCID: PMCPMC7319546.

30. Wen B, Wang X, Zhang B. PepQuery enables fast, accurate, and convenient proteomic validation of novel genomic alterations. Genome Res. 2019;29(3):485–93. Epub 20190104. doi: 10.1101/gr.235028.118. PubMed PMID: 30610011; PubMed Central PMCID: PMCPMC6396417.

31. Camacho C, Coulouris G, Avagyan V, Ma N, Papadopoulos J, Bealer K, et al. BLAST+: architecture and applications. BMC Bioinformatics. 2009;10:421. Epub 20091215. doi: 10.1186/1471-2105-10-421. PubMed PMID: 20003500; PubMed Central PMCID: PMCPMC2803857.

32. Cimen Bozkus C, Blazquez AB, Enokida T, Bhardwaj N. A T-cell-based immunogenicity protocol for evaluating human antigen-specific responses. STAR Protoc. 2021;2(3):100758. Epub 20210817. doi: 10.1016/j.xpro.2021.100758. PubMed PMID: 34458873; PubMed Central PMCID: PMCPMC8377590.

33. Vita R, Blazeska N, Marrama D, Members ICT, Duesing S, Bennett J, et al. The Immune Epitope Database (IEDB): 2024 update. Nucleic Acids Res. 2025;53(D1):D436-D43. doi: 10.1093/nar/gkae1092. PubMed PMID: 39558162; PubMed Central PMCID: PMCPMC11701597.

34. Available from: www.iedb.org.

35. Li Y, Zhang Y, Pan T, Zhou P, Zhou W, Gao Y, et al. Shedding light on the hidden human proteome expands immunopeptidome in cancer. Brief Bioinform. 2022;23(2). doi: 10.1093/bib/bbac034. PubMed PMID: 35189633.

36. Prensner JR, Abelin JG, Kok LW, Clauser KR, Mudge JM, Ruiz-Orera J, et al. What Can Ribo-Seq, Immunopeptidomics, and Proteomics Tell Us About the Noncanonical Proteome? Mol Cell Proteomics. 2023;22(9):100631. Epub 20230811. doi: 10.1016/j.mcpro.2023.100631. PubMed PMID: 37572790; PubMed Central PMCID: PMCPMC10506109.

37. Vita R, Mahajan S, Overton JA, Dhanda SK, Martini S, Cantrell JR, et al. The Immune Epitope Database (IEDB): 2018 update. Nucleic Acids Res. 2019;47(D1):D339-D43. doi: 10.1093/nar/gky1006. PubMed PMID: 30357391; PubMed Central PMCID: PMCPMC6324067.

38. Gonzalez-Galarza FF, McCabe A, Santos E, Jones J, Takeshita L, Ortega-Rivera ND, et al. Allele frequency net database (AFND) 2020 update: gold-standard data classification, open access genotype data and new query tools. Nucleic Acids Res. 2020;48(D1):D783–D8. doi: 10.1093/nar/gkz1029. PubMed PMID: 31722398; PubMed Central PMCID: PMCPMC7145554.

39. Satapornpong P, Jinda P, Jantararoungtong T, Koomdee N, Chaichan C, Pratoomwun J, et al. Genetic Diversity of HLA Class I and Class II Alleles in Thai Populations: Contribution to Genotype-Guided Therapeutics. Front Pharmacol. 2020;11:78. Epub 20200227. doi: 10.3389/fphar.2020.00078. PubMed PMID: 32180714; PubMed Central PMCID: PMCPMC7057685.

40. Nurul-Aain AF, Tan LK, Heselynn H, Nor-Shuhaila S, Eashwary M, Wahinuddin S, et al. HLA-A, -B, -C, -DRB1 and -DQB1 alleles and haplotypes in 271 Southeast Asia Indians from Peninsular Malaysia. Hum Immunol. 2020;81(6):263–4. Epub 20200418. doi: 10.1016/j.humimm.2020.04.004. PubMed PMID: 32312605.

41. Chandanayingyong D, Stephens HA, Klaythong R, Sirikong M, Udee S, Longta P, et al. HLA-A, -B, -DRB1, -DQA1, and -DQB1 polymorphism in Thais. Hum Immunol. 1997;53(2):174-82. doi: 10.1016/S0198-8859(96)00284-4. PubMed PMID: 9129976.

42. Chong C, Muller M, Pak H, Harnett D, Huber F, Grun D, et al. Integrated proteogenomic deep sequencing and analytics accurately identify non-canonical peptides in tumor immunopeptidomes. Nat Commun. 2020;11(1):1293. Epub 20200310. doi: 10.1038/s41467-020-14968-9. PubMed PMID: 32157095; PubMed Central PMCID: PMCPMC7064602.

43. Sun S, Liu L, Zhang J, Sun L, Shu W, Yang Z, et al. The role of neoantigens and tumor mutational burden in cancer immunotherapy: advances, mechanisms, and perspectives. J Hematol Oncol. 2025;18(1):84. Epub 20250902. doi: 10.1186/s13045-025-01732-z. PubMed PMID: 40898324; PubMed Central PMCID: PMCPMC12406617.

44. Sholl LM, Hirsch FR, Hwang D, Botling J, Lopez-Rios F, Bubendorf L, et al. The Promises and Challenges of Tumor Mutation Burden as an Immunotherapy Biomarker: A Perspective from the International Association for the Study of Lung Cancer Pathology Committee. J Thorac Oncol. 2020;15(9):1409–24. Epub 20200606. doi: 10.1016/j.jtho.2020.05.019. PubMed PMID: 32522712; PubMed Central PMCID: PMCPMC8363213.

45. Amber Miller P, Laura Cattaneo, Yan W Asmann, PhD, Esteban Braggio, PhD, Jonathan J Keats, PhD, Daniel Auclair, PhD, Sagar Lonial, MD FACP, The MMRF CoMMpass Network, Stephen J. Russell, MD PhD, A. Keith Stewart, MD. Patient-Specific Mutation-Derived Tumor Antigens As Targets for Cancer Immunotherapy in Multiple Myeloma. Blood 2016;126(23). doi: 10.1182/blood.V126.23.1851.1851.

46. Panchenko MV. Structure, function and regulation of jade family PHD finger 1 (JADE1). Gene. 2016;589(1):1–11. Epub 20160504. doi: 10.1016/j.gene.2016.05.002. PubMed PMID: 27155521; PubMed Central PMCID: PMCPMC4903948.

47. Foy RL, Song IY, Chitalia VC, Cohen HT, Saksouk N, Cayrou C, et al. Role of Jade-1 in the histone acetyltransferase (HAT) HBO1 complex. J Biol Chem. 2008;283(43):28817–26. Epub 20080806. doi: 10.1074/jbc.M801407200. PubMed PMID: 18684714; PubMed Central PMCID: PMCPMC2570895.

48. Yadav M, Jhunjhunwala S, Phung QT, Lupardus P, Tanguay J, Bumbaca S, et al. Predicting immunogenic tumour mutations by combining mass spectrometry and exome sequencing. Nature. 2014;515(7528):572–6. doi: 10.1038/nature14001. PubMed PMID: 25428506.

49. Darshit shah AR-W, Ankur Dhanik, Kamil Cygan, Olav Olsen, William Olson, Robert Salzler. Proteogenomics and de novo sequencing based approach for neoantigen discovery from the immunopeptidomes of patient CRC liver metastases using Mass Spectrometry. The Journal of Immunology. 2020;204(1_Supplement). doi: 10.4049/jimmunol.204.Supp.217.16.

