## Supplementary figures and images for "Mass Spectrometry-Based Profiling of Personalized Immunopeptidomes in Thai Renal Cell Carcinoma"

### Fig S1

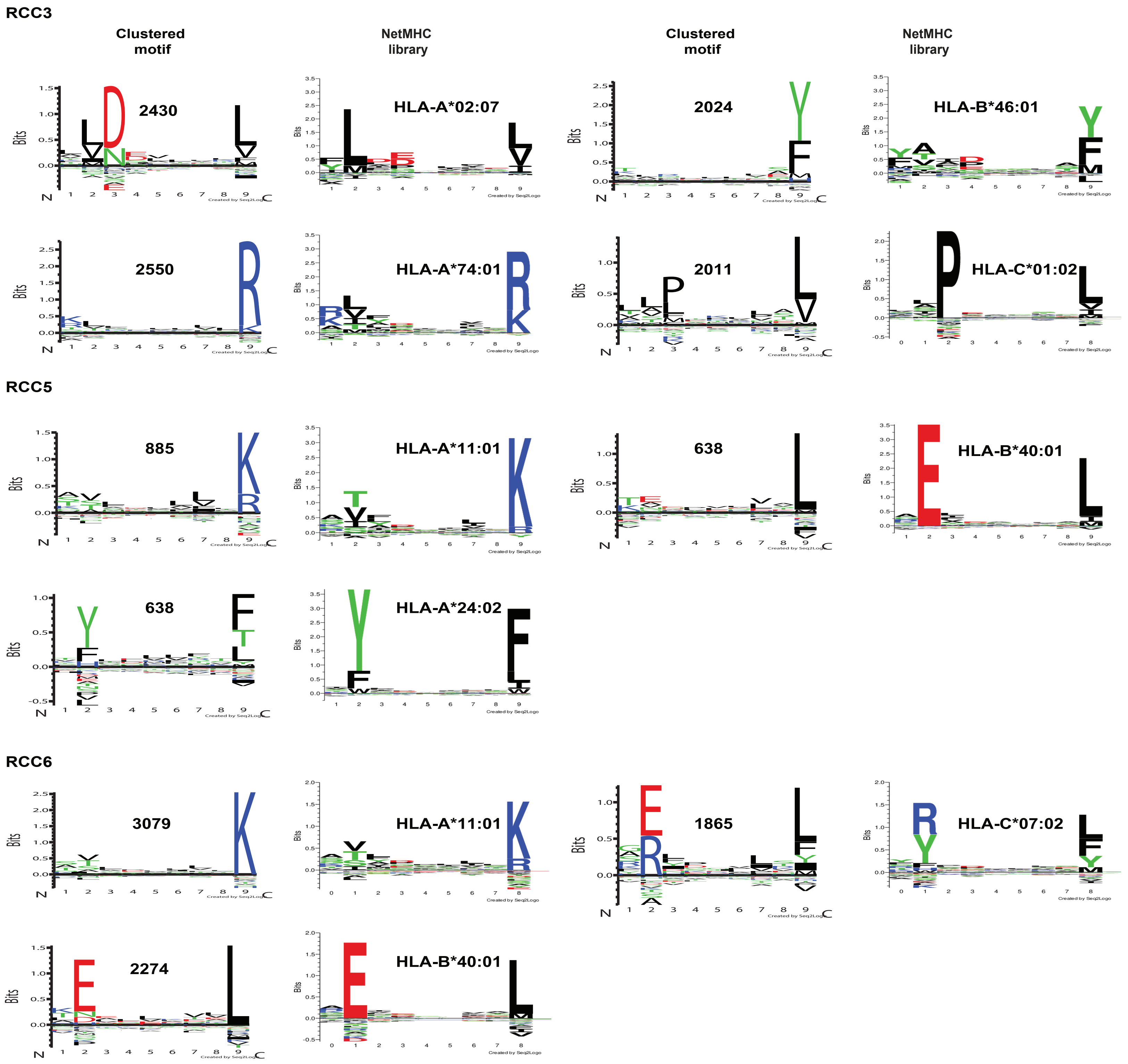

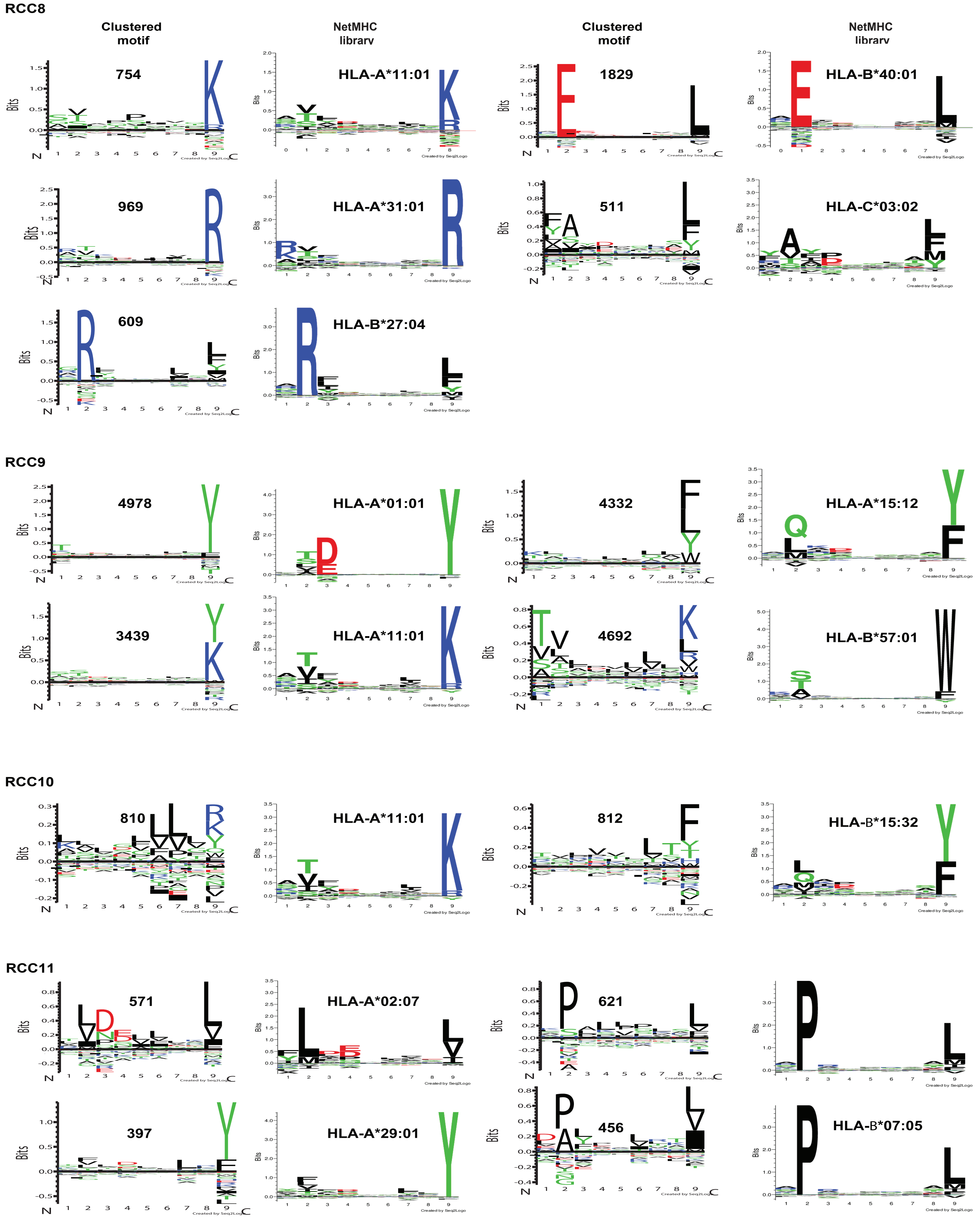

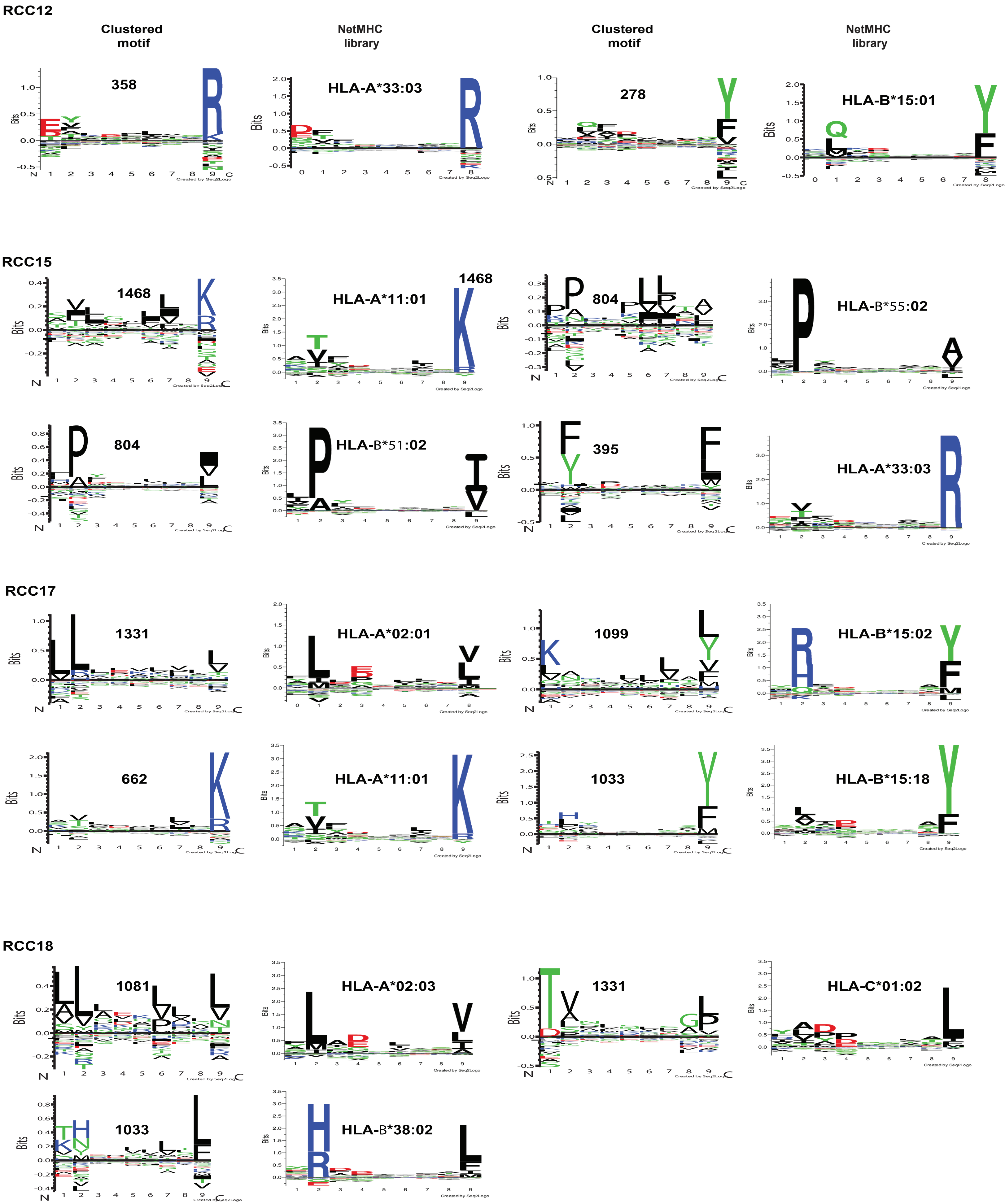

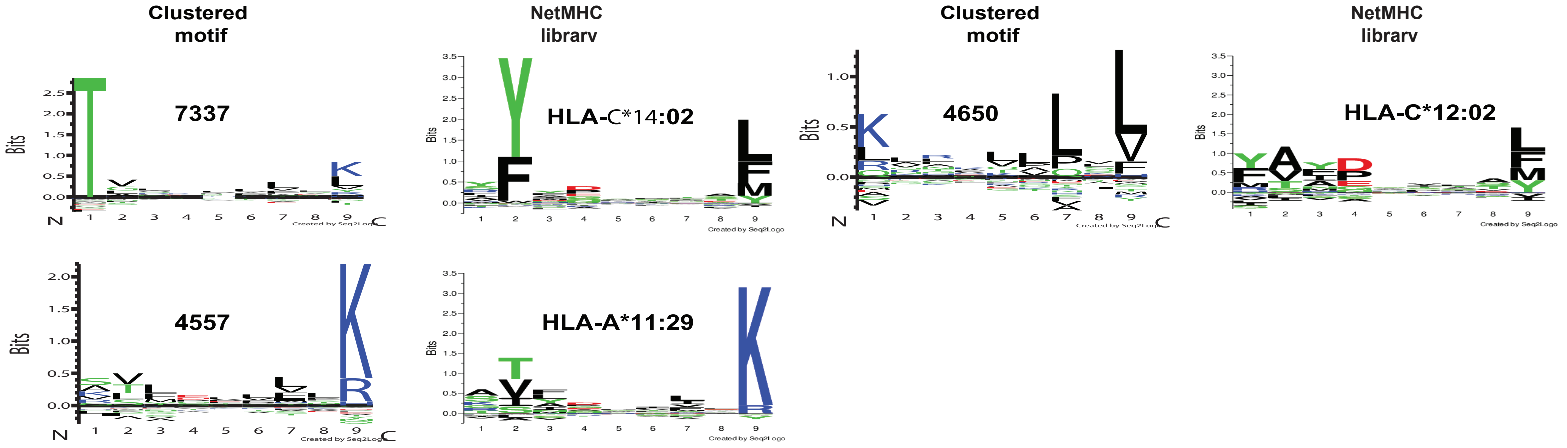
