## Supplementary material for "Mass Spectrometry-Based Profiling of Personalized Immunopeptidomes in Thai Renal Cell Carcinoma": Fig S2

CU01

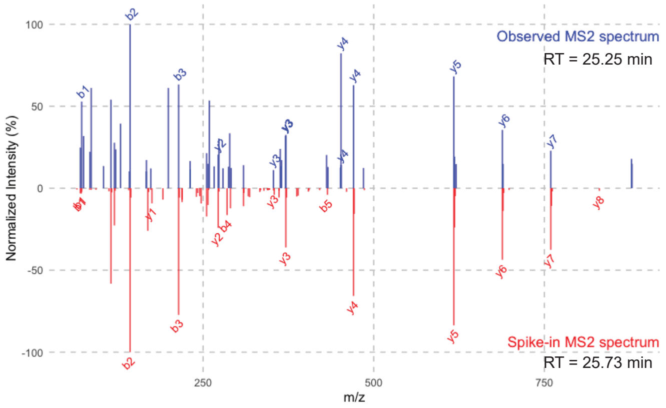


CU02

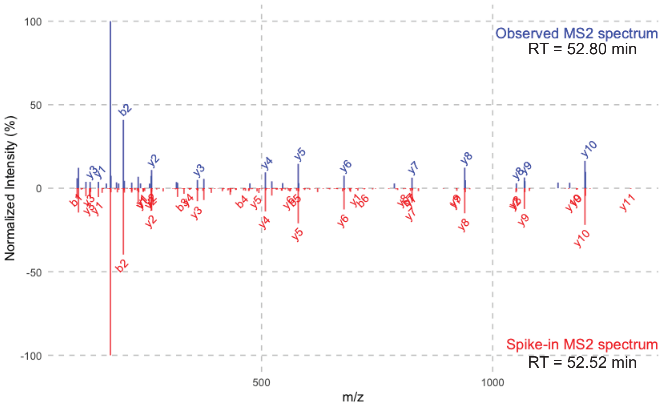


CU03

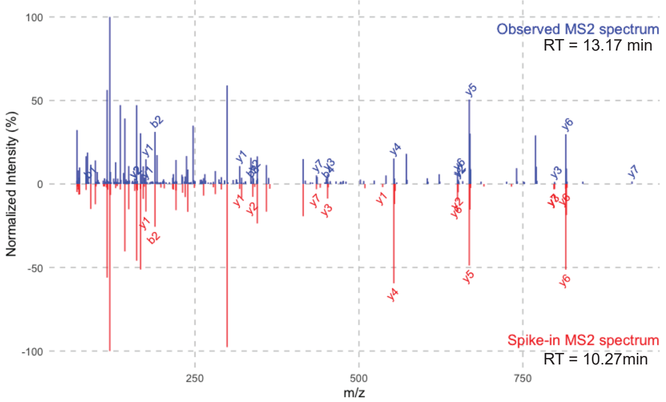


CU04

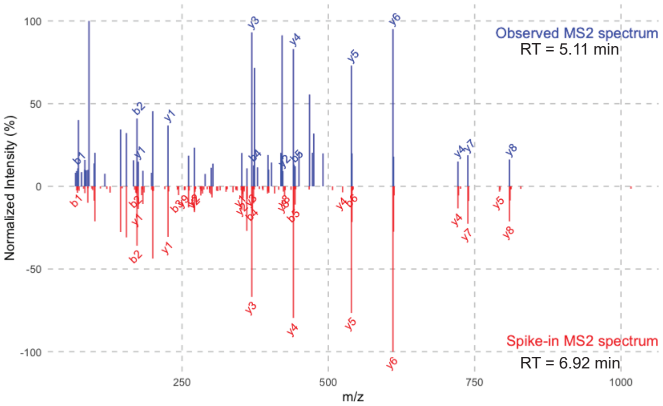


CU05

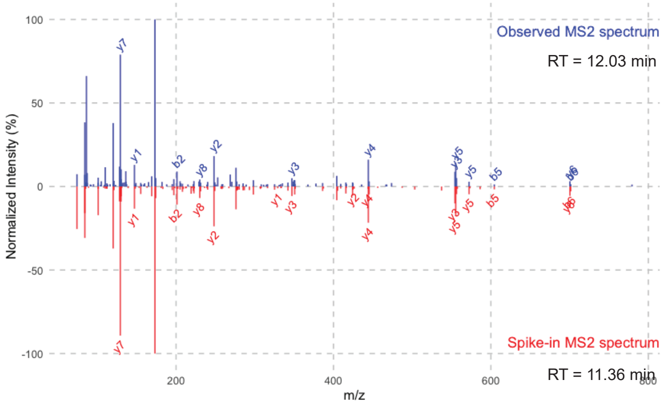


CU06

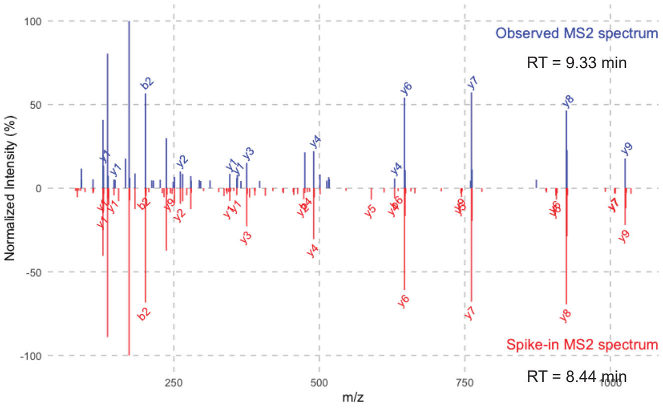


CU07

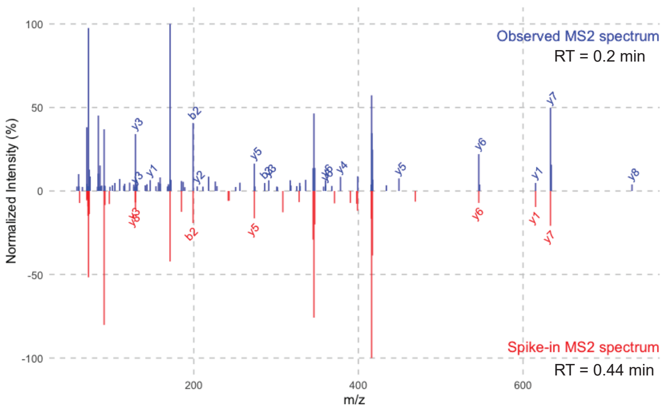


CU08

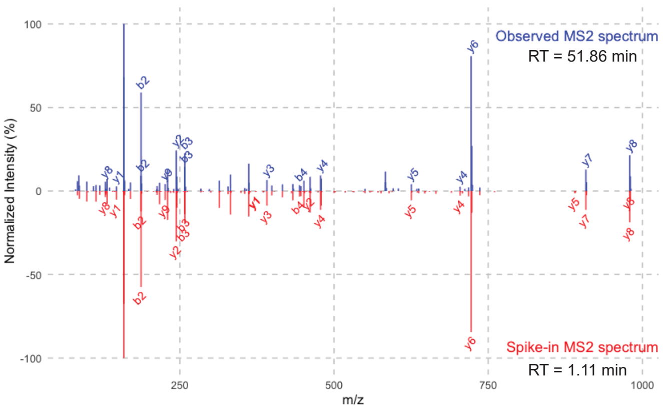

CU09

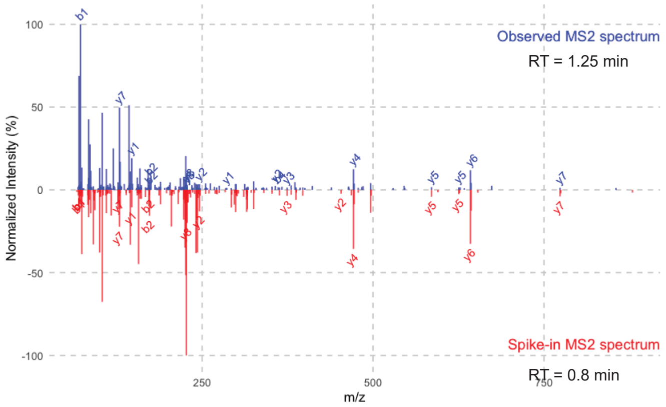


CU10

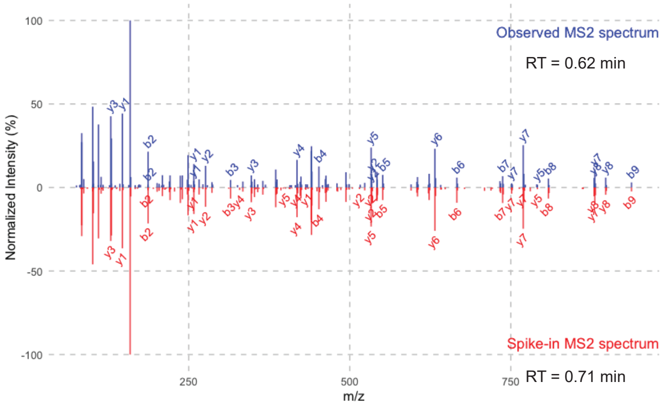


CU11

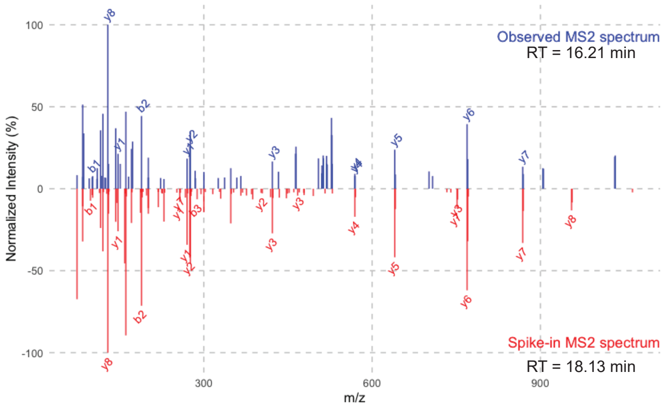


CU13

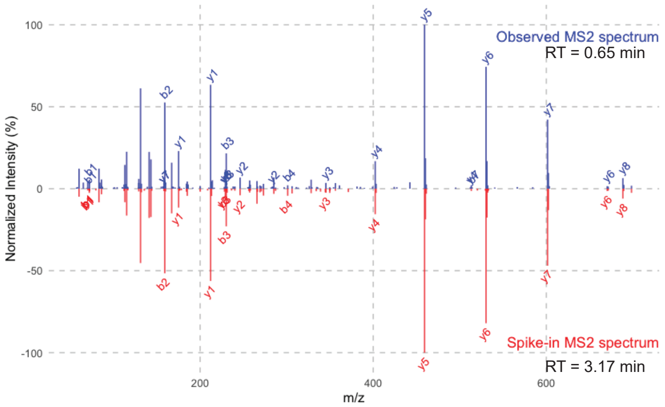


CU15

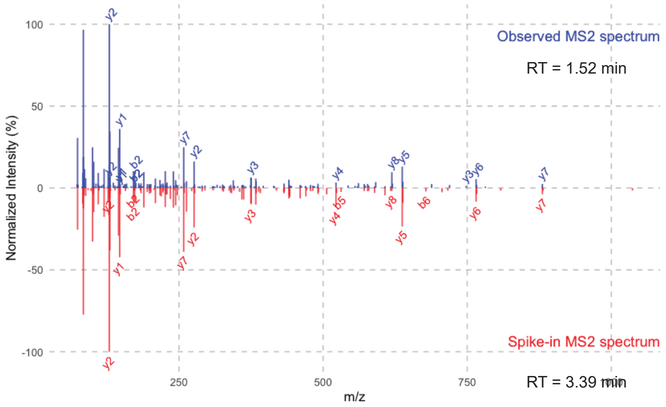


CU16

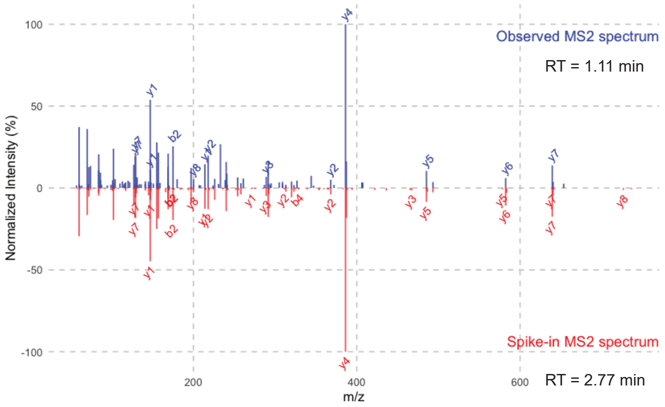


CU17

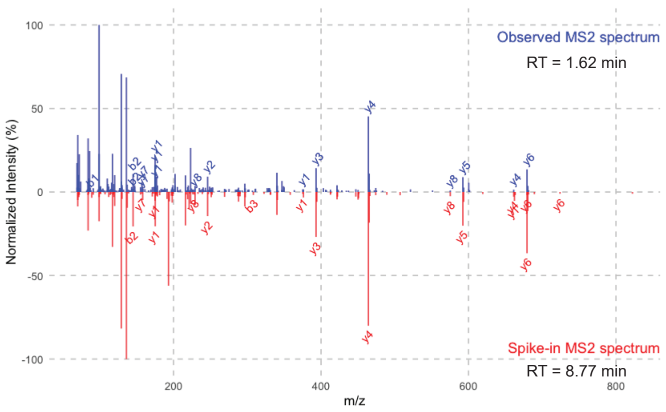


CU19

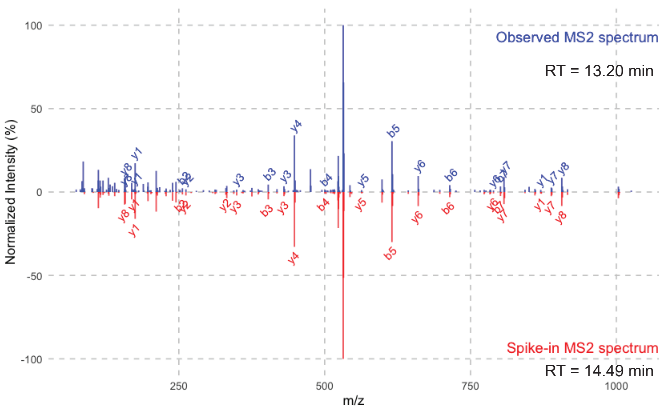

CU21

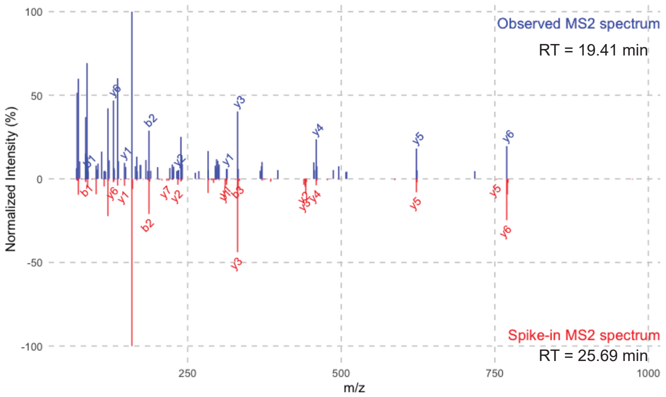


CU22

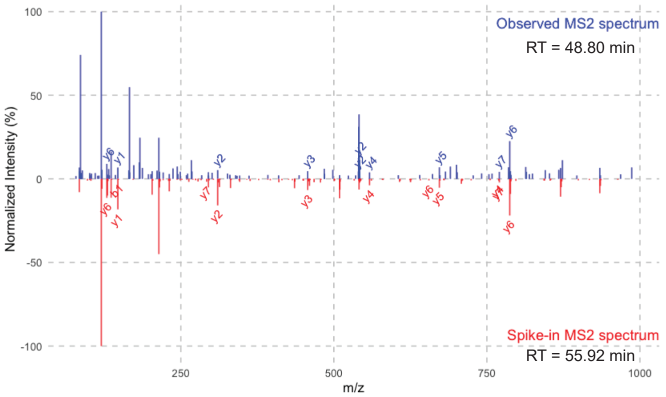


CU23

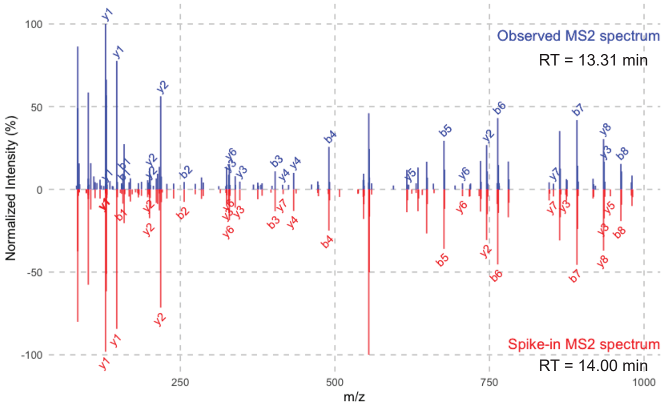


CU24

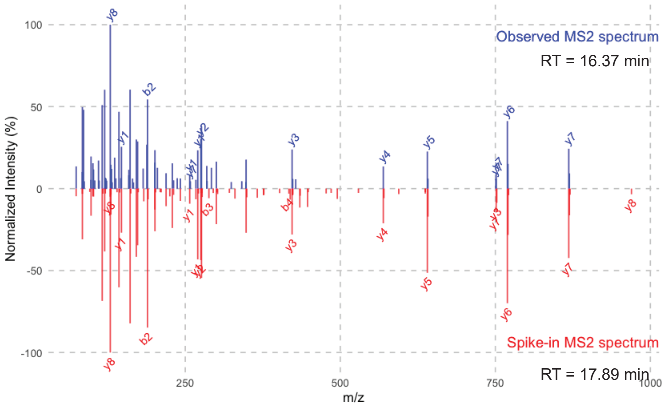


CU25

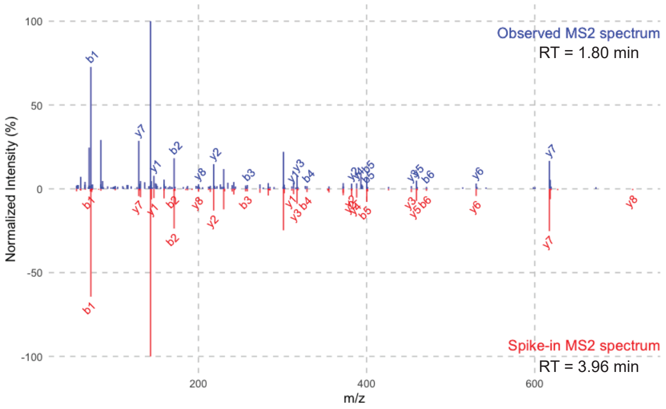


CU27

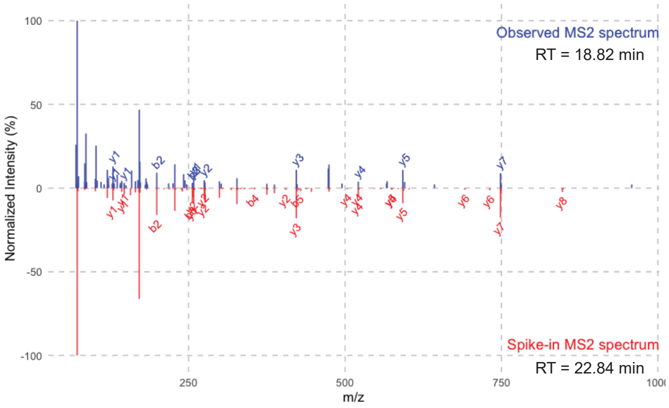


CU28

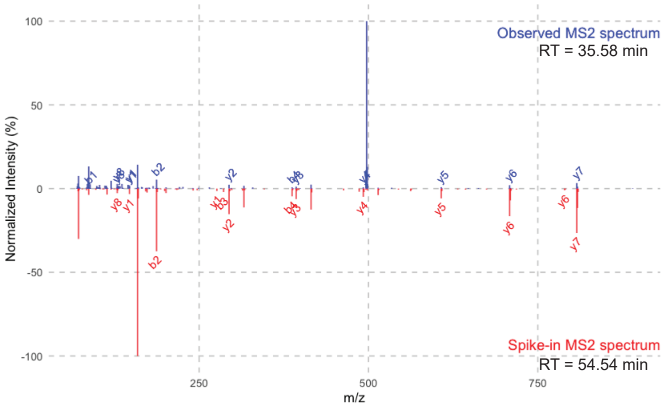


CU30

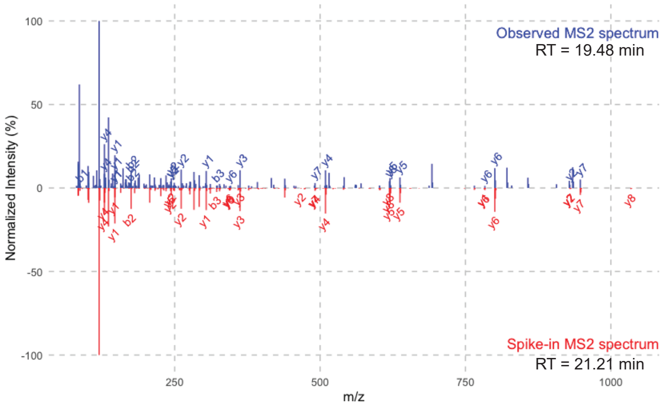

CU31

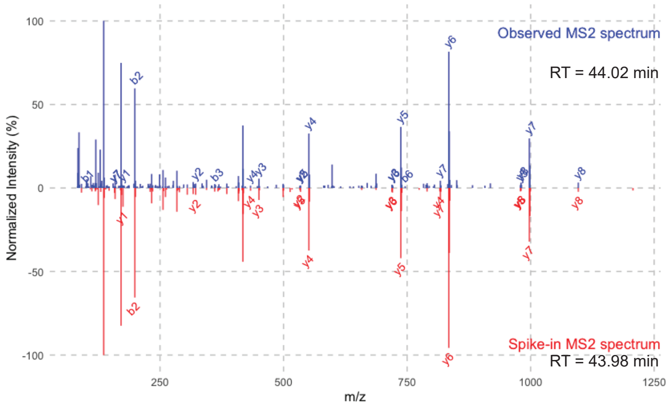


CU32

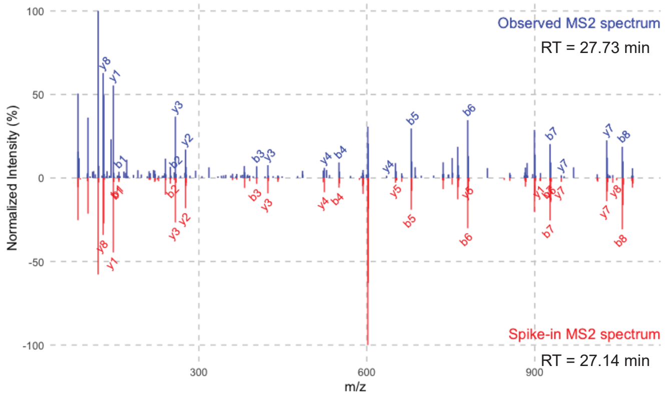


CU33

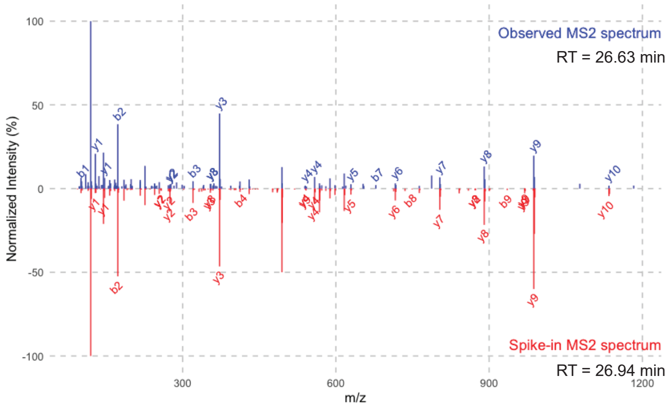


CU35

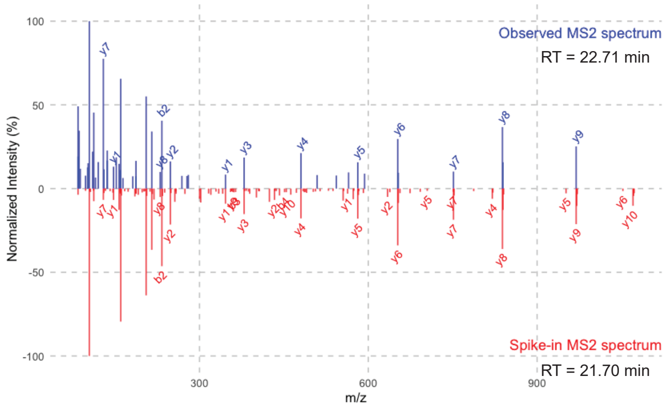


CU36

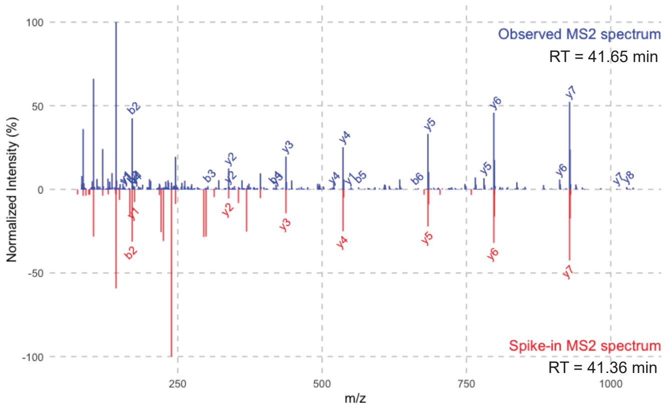


CU37

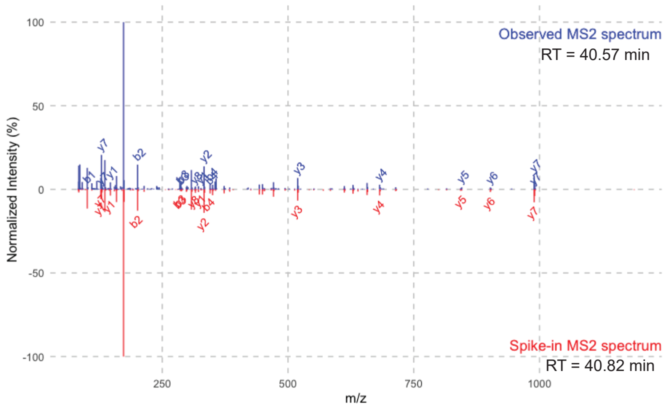


CU38

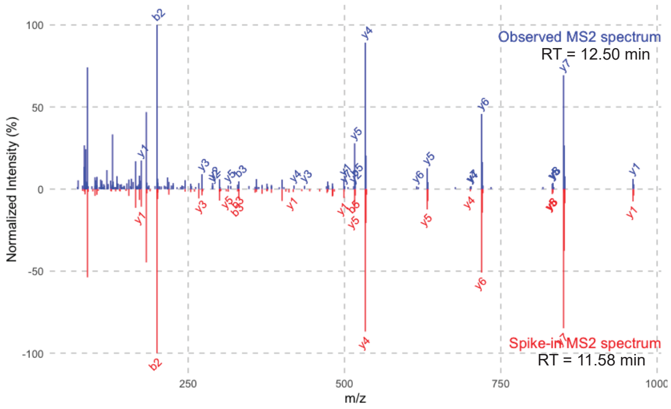


CU39

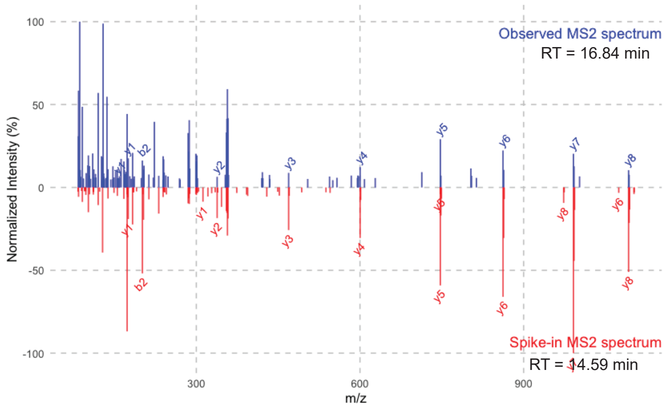

CU40

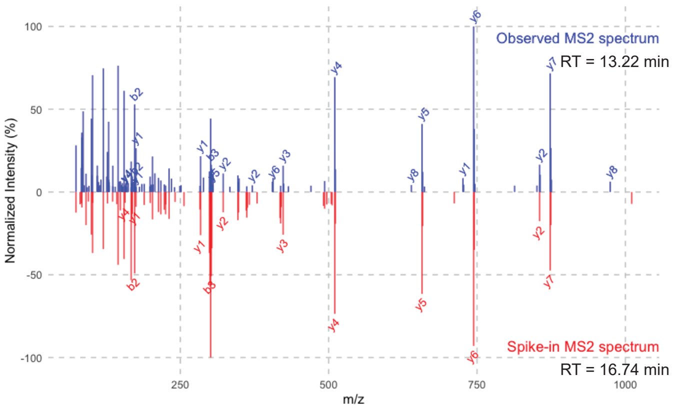


CU41

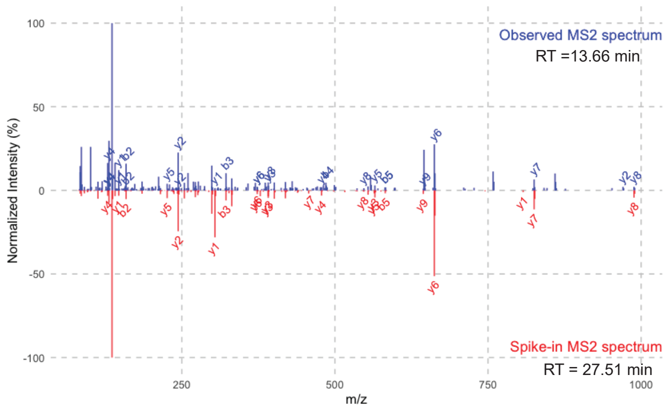


CU42

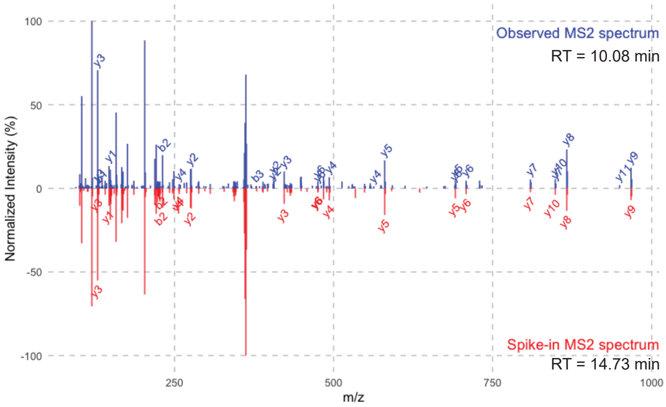


CU43

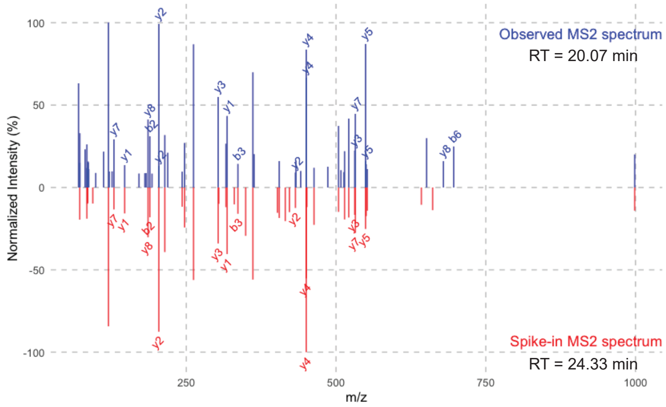


CU44

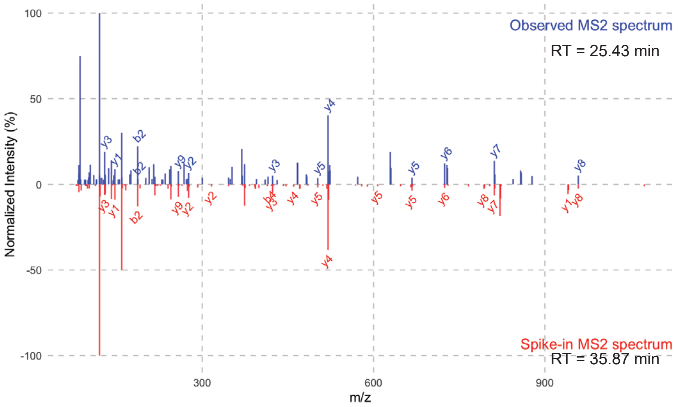


CU45

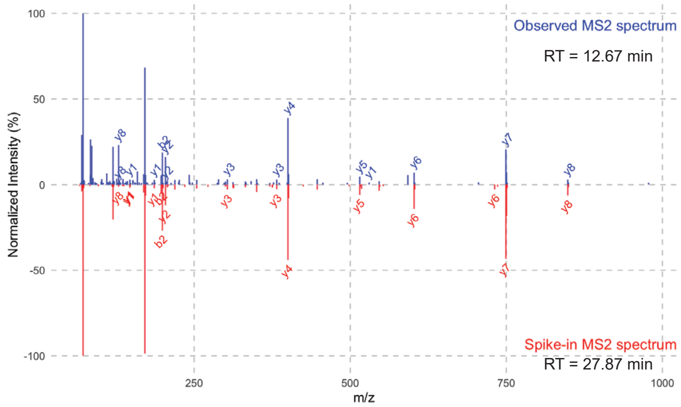


CU46

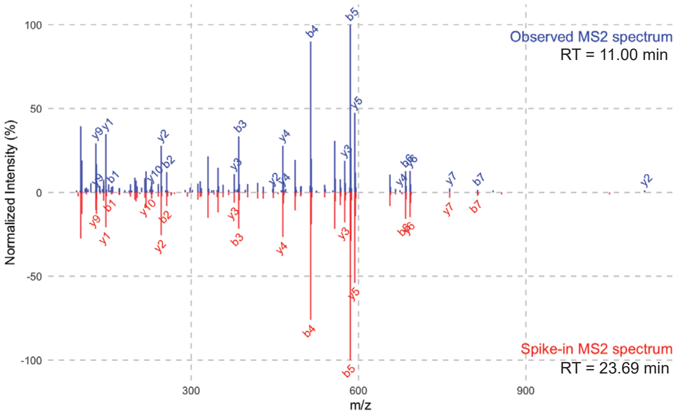


CU47

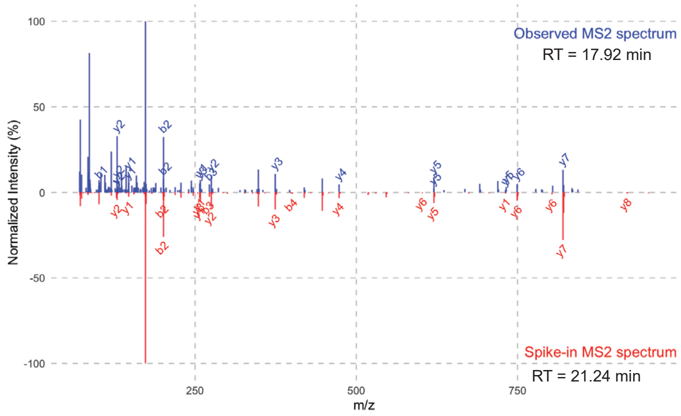

CU48

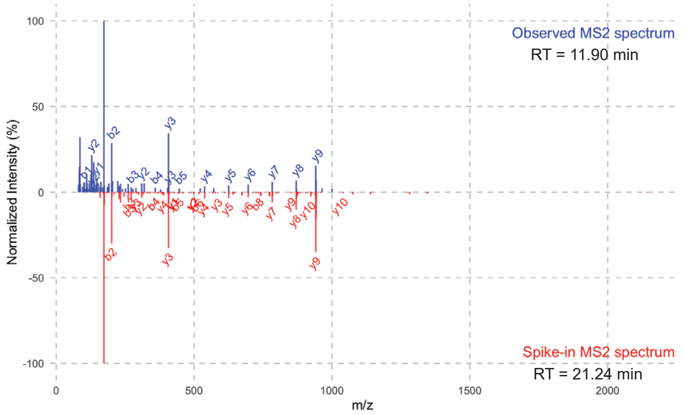


CU49

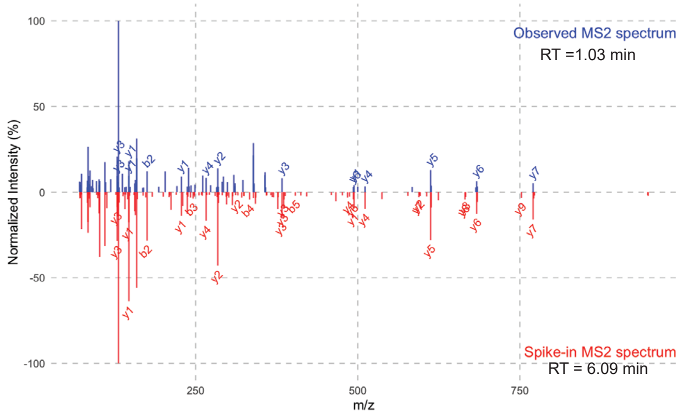


CU50

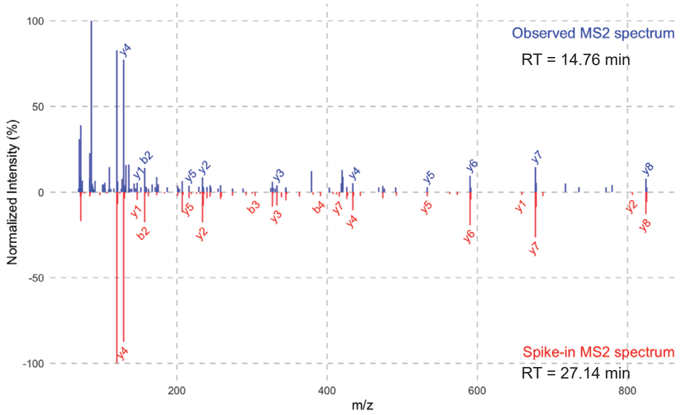


CU51

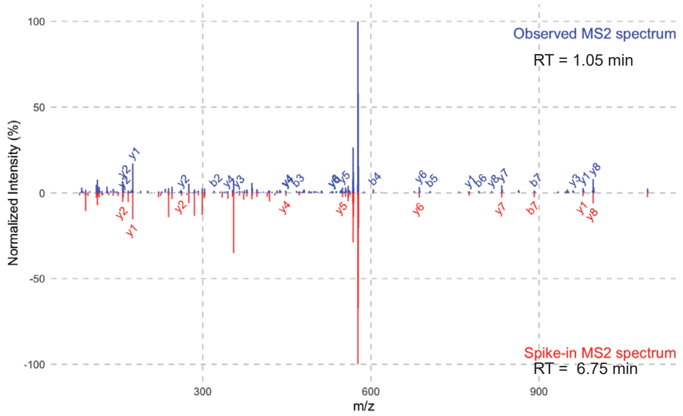


CU52

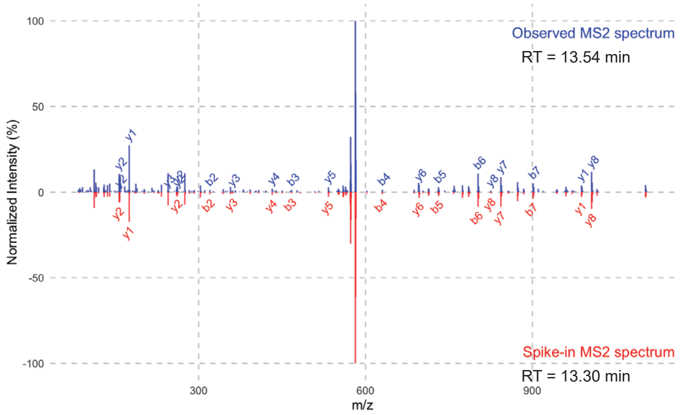


CU53

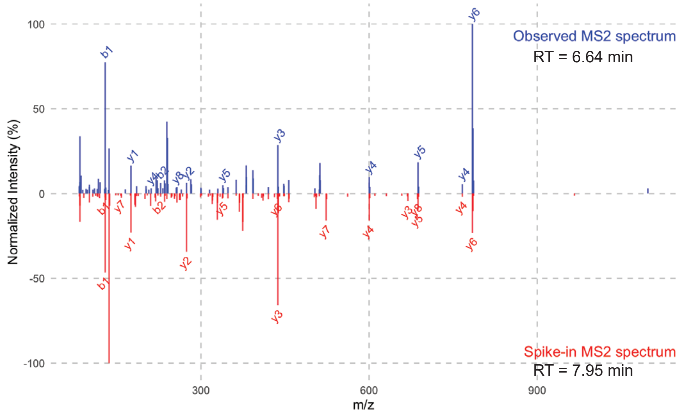


CU54

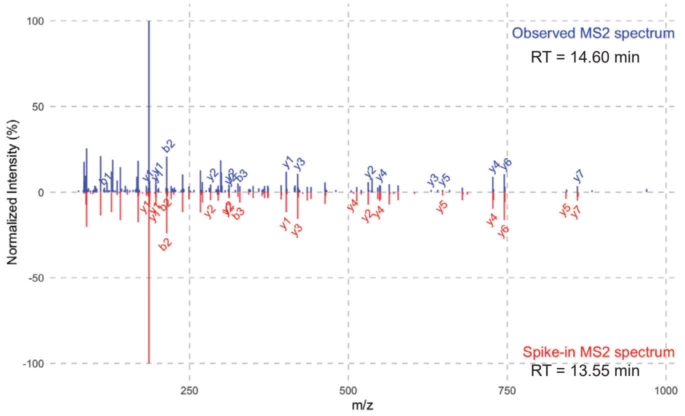


CU56

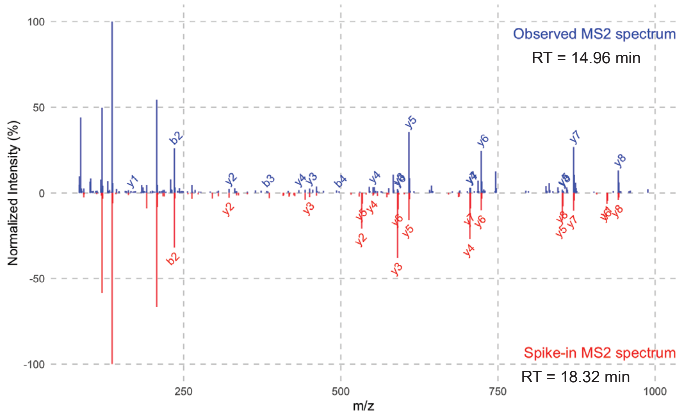
